# SARS-CoV-2 ferritin nanoparticle vaccines elicit broad SARS coronavirus immunogenicity

**DOI:** 10.1101/2021.05.09.443331

**Authors:** M. Gordon Joyce, Wei-Hung Chen, Rajeshwer S. Sankhala, Agnes Hajduczki, Paul V. Thomas, Misook Choe, William Chang, Caroline E. Peterson, Elizabeth Martinez, Elaine B. Morrison, Clayton Smith, Aslaa Ahmed, Lindsay Wieczorek, Alexander Anderson, Rita E. Chen, James Brett Case, Yifan Li, Therese Oertel, Lorean Rosado, Akshaya Ganesh, Connor Whalen, Joshua M. Carmen, Letzibeth Mendez-Rivera, Christopher Karch, Neelakshi Gohain, Zuzana Villar, David McCurdy, Zoltan Beck, Jiae Kim, Shikha Shrivastava, Ousman Jobe, Vincent Dussupt, Sebastian Molnar, Ursula Tran, Chandrika B. Kannadka, Michelle Zemil, Htet Khanh, Weimin Wu, Matthew A. Cole, Debra K. Duso, Larry W. Kummer, Tricia J. Lang, Shania E. Muncil, Jeffrey R. Currier, Shelly J. Krebs, Victoria R. Polonis, Saravanan Rajan, Patrick M. McTamney, Mark T. Esser, William W. Reiley, Morgane Rolland, Natalia de Val, Michael S. Diamond, Gregory D. Gromowski, Gary R. Matyas, Mangala Rao, Nelson L. Michael, Kayvon Modjarrad

## Abstract

The need for SARS-CoV-2 next-generation vaccines has been highlighted by the rise of variants of concern (VoC) and the long-term threat of other coronaviruses. Here, we designed and characterized four categories of engineered nanoparticle immunogens that recapitulate the structural and antigenic properties of prefusion Spike (S), S1 and RBD. These immunogens induced robust S-binding, ACE2-inhibition, and authentic and pseudovirus neutralizing antibodies against SARS-CoV-2 in mice. A Spike-ferritin nanoparticle (SpFN) vaccine elicited neutralizing titers more than 20-fold higher than convalescent donor serum, following a single immunization, while RBD-Ferritin nanoparticle (RFN) immunogens elicited similar responses after two immunizations. Passive transfer of IgG purified from SpFN- or RFN-immunized mice protected K18-hACE2 transgenic mice from a lethal SARS-CoV-2 virus challenge. Furthermore, SpFN- and RFN-immunization elicited ACE2 blocking activity and neutralizing ID50 antibody titers >2,000 against SARS-CoV-1, along with high magnitude neutralizing titers against major VoC. These results provide design strategies for pan-coronavirus vaccine development.

**HIGHLIGHTS:**

- Iterative structure-based design of four Spike-domain Ferritin nanoparticle classes of immunogens
- SpFN-ALFQ and RFN-ALFQ immunization elicits potent neutralizing activity against SARS-CoV-2, variants of concern, and SARS-CoV-1
- Passively transferred IgG from immunized C57BL/6 mice protects K18-hACE2 mice from lethal SARS-CoV-2 challenge

## INTRODUCTION

Seven coronaviruses (CoV) cause disease in humans, with three of these, SARS-CoV-1, MERS-CoV, and SARS-CoV-2 having emerged since 2003 (Cui et al., 2019) and displaying high mortality rates. SARS-CoV-2 is easily transmitted by humans and created a pandemic, infecting over 100 million people, causing over 2 million deaths to date, and resulting in an urgent need for protective and durable vaccines. Rapid vaccine development in a worldwide effort, led to the evaluation of hundreds of SARS-CoV-2 vaccine candidates and rapid worldwide vaccine distribution and use.

The response to SARS-CoV-2 was facilitated by multiple efforts over the last decade to enable CoV pandemic preparedness, initially based on MERS-CoV vaccine design and development (Wang et al., 2015), phase I vaccine trials (Modjarrad et al., 2019), and a global effort by the Coalition for Epidemic Preparedness Innovations (CEPI) to advance vaccine candidates (Plotkin, 2017). The elucidation of CoV Spike (S) glycoprotein structures (Kirchdoerfer et al., 2016; Walls et al., 2016) allowed structure-based vaccine design of stabilized S glycoprotein immunogens from multiple CoVs (Pallesen et al., 2017), providing a blueprint for SARS-CoV-2 vaccine design (Corbett et al., 2020).

The CoV S protein mediates virus entry, is immunogenic (Iyer et al., 2020; Wang et al., 2021) and encodes multiple neutralizing epitopes (Greaney et al., 2021) making it the primary target for natural and vaccine-induced CoV humoral immunity and vaccine design (Jiang et al., 2020) and the target of most COVID-19 vaccines. S is a class I fusion glycoprotein consisting of a S1 attachment subunit and S2 fusion subunit that remain non-covalently associated in a metastable, heterotrimeric S on the virion surface (Walls et al., 2020). In the S1 subunit, there is a N-terminal domain (NTD), and C-terminal domain (CTD) that includes the receptor-binding domain (RBD). The RBD binds to the human angiotensin converting enzyme 2 (hACE2) facilitating cell entry (Lan et al., 2020). Multiple antigenic sites have been identified on the S protein, including distinct sites on the RBD and the S1 domain, including an NTD supersite (Brouwer et al., 2020; Cerutti et al., 2021; Liu et al., 2020; Zost et al., 2020). Convalescent serum antibodies or monoclonal antibodies capable of potently inhibiting infection *in vitro* can reduce disease severity or mortality in rodents, non-human primates (Barnes et al., 2020) and humans (Duan et al., 2020; Salazar et al., 2020; Shen et al., 2020).

Due to the unknown parameters of SARS-CoV-2 vaccine durability, specific age- or population-needs, emergence of SARS-CoV-2 variants of concern (VoC) (Wibmer et al., 2020), and the constant threat of emerging CoV pathogens (Menachery et al., 2015), second-generation COVID-19 or pan-sarbecovirus vaccines will be needed. Iterative structure-based design for viral glycoproteins (McLellan et al., 2013; Joyce et al., 2016) stabilizing neutralizing epitopes or epitope-based vaccine design (Chen et al., 2021; Kong et al., 2019) show that rational vaccine design can lead to the elicitation of broad immune responses. Broad cross-reactive responses elicited by engineered vaccines have also been advanced for influenza (Boyoglu-Barnum et al., 2020; Kanekiyo et al., 2019). In the case of CoVs, a set of cross-reactive epitopes have recently been described (Barnes et al., 2020; Joyce et al., 2020; Sauer et al., 2021; Wrapp et al., 2020), with many of the preferred neutralizing antibodies centered on the RBD domain (Li et al., 2021; Pinto et al., 2020; Rappazzo et al., 2021).

Next-generation strategies to augment specific immune responses as well as enhance cross-reactivity include the use of nanoparticle vaccine technology (Cohen et al., 2021) and next-generation adjuvants. Nanoparticle technologies have been shown to improve antigen structure and stability, as well as vaccine targeted delivery, immunogenicity, with good safety profiles (Pati et al., 2018). Engineered nanoparticle-vaccines can elicit broader immune responses (Darricarrere et al., 2021; Kanekiyo et al., 2019; Kanekiyo et al., 2013) or more efficacious immune responses (Kanekiyo et al., 2015). The repetitive array of the viral surface component allows for robust B-cell activation facilitating memory B cell expansion and generation of long-lived plasma cells. More recently, in efforts to generate more effective vaccines that can prevent infection by resistant pathogens such as HIV-1 or Influenza, a set of engineered nanoparticle vaccines have been developed. Utilizing naturally occurring nanoparticle molecules such as bacterial ferritin, antigens are fused to the ferritin molecule to recapitulate complex trimeric class I glycoproteins, and to increase the immune response for weakly immunogenic targets. Nanoparticle technologies have also been shown to improve antigen structure and stability, as well as vaccine targeted delivery, immunogenicity and safety (Pati et al., 2018). Recently designed single and multi-component nanoparticle vaccines (Brouwer et al., 2021; Walls et al., 2020) show promise from both an immunological (Cawlfield et al., 2019; Marcandalli et al., 2019) and a cGMP production perspective (Ueda et al., 2020).

Engineered nanoparticle vaccines and their capacity to generate enhanced immune responses in humans are currently being studied and include influenza (NCT03186781; NCT03814720; NCT04579250), Epstein-Barr virus (NCT04645147), malaria (NCT04296279) and a recently described SARS-CoV-2 nanoparticle vaccine (IVX-411) (Walls et al., 2020). Use of potent adjuvants such as liposomal-saponin adjuvants can further enhance the protective immune response (Cawlfield et al., 2019; Lal et al., 2015; Om et al., 2020) even in the context of nanoparticle vaccines (Langowski et al., 2020) (Kaba et al., 2018). Based on the results described herein and data from associated non-human primate experiments (Joyce et al., 2021; King et al., 2021), a S-Ferritin immunogen with a liposomal adjuvant, ALFQ is currently being assessed in a phase I clinical trial (NCT04784767).

Here we report the structure-based design and pre-clinical assessment of four categories of S-domain ferritin nanoparticles including stabilized S-trimer-ferritin nanoparticles (SpFN), RBD-ferritin nanoparticles (RFN), S1-ferritin nanoparticles, and RBD-NTD-ferritin nanoparticles. By using a set of biophysical, structural, and antigenic assessments, combined with animal immunogenicity testing, we identify multiple immunogens that elicit substantial neutralizing antibody titers against SARS-CoV-2 and related VoC. These antibody levels provide robust protection against SARS-CoV-2 challenge in the K18-hACE2 mouse model. We further show that subsequent immunizations not only increase the SARS-CoV-2 neutralization titer, but also expand the neutralization breadth and titer against the heterologous SARS-CoV-1 virus. These data provide multiple immunogen design strategies for pan-betacoronavirus vaccine development.

## RESULTS

### Immunogen Design of SARS-CoV-2 S-domain Ferritin Nanoparticles

Using the initial SARS-CoV-2 genome sequence (Genbank: MN9089473), we designed four categories of S-domain ferritin-fusion recombinant proteins as immunogens for expression as nanoparticles based on the major antigenic domains of the S ectodomain (Figure S1). The *Helicobacter pylori* ferritin molecule was genetically linked to the C-terminal region of the following S antigens (i) S ectodomain (residues 12-1158) (ii) RBD (residues 331-527), (iii) RBD linked in tandem to the NTD (residues 12-303), and (iv) S1 (residues 12-696) (Figure 1, Figure S1, and Table S1). In the case of the Spike ferritin nanoparticle designs, a short linker to the ferritin molecule was used to utilize the natural three-fold axis, for display of eight Spikes. In the case of the other designs, a short region of bullfrog ferritin was utilized to allow equidistant distribution of the 24 S-domain molecules on the ferritin surface (Figure 1). Our overall approach was to compare and contrast the immunogen structures, antigenicity, immunogenicity elicited by these different immunogens with the goal to identify the best immunogen to take forward into further development.

**Figure 1.**
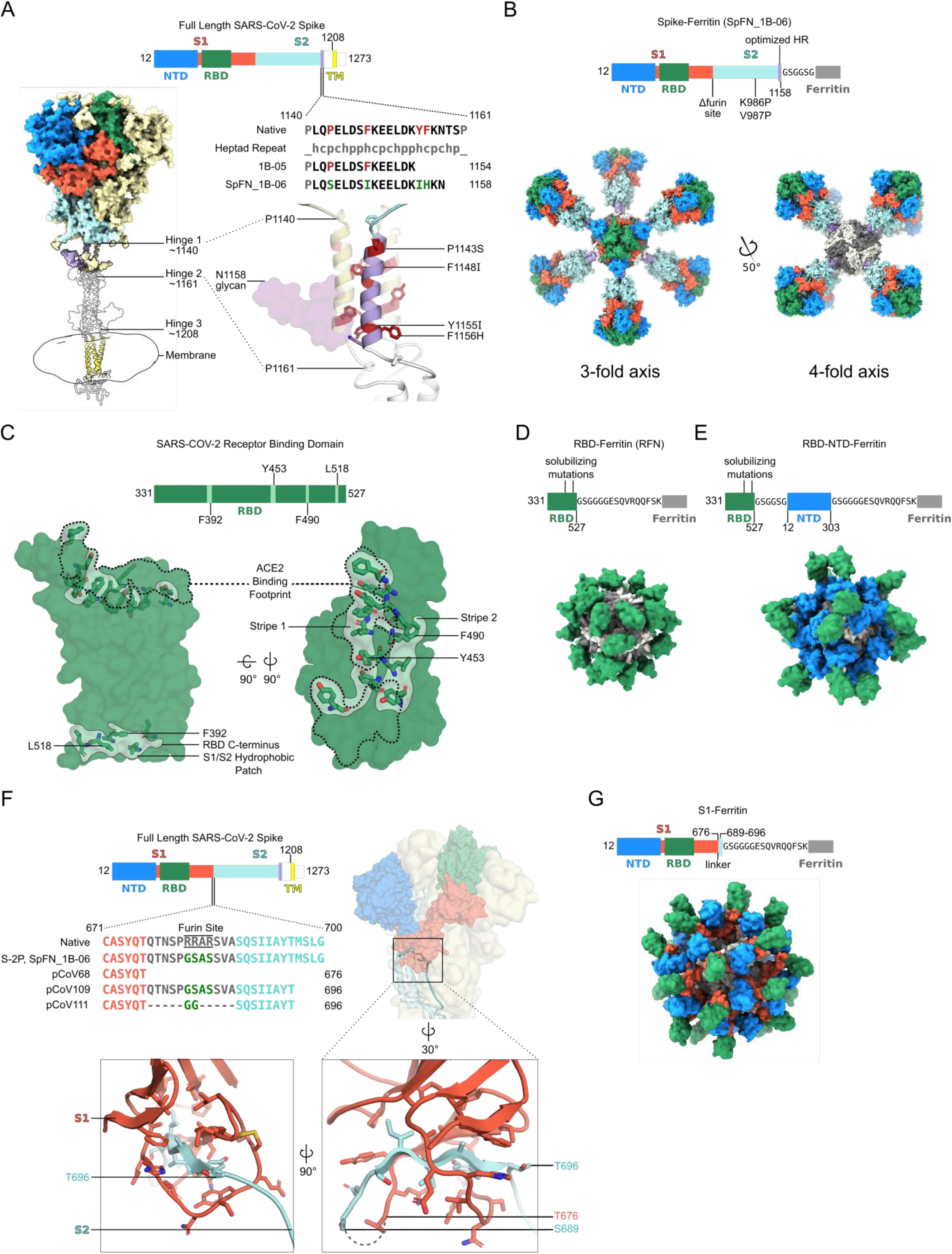
Structure-based design of SARS-CoV-2 S-based ferritin nanoparticle immunogens. (A) Full length SARS-CoV-2 S schematic and 3D-structure. S hinges identified by molecular dynamics simulations and electron cryotomography are labeled on the 3D-model ((Turonova et al., 2020). The structured trimeric ectodomain is colored according to the schematic with the N-terminal domain (NTD) and Receptor-Binding Domain (RBD) of the S1 polypeptide and the C-terminal coiled coil N-terminal to hinge 1 colored blue, green, and purple, respectively. Remaining portions of the S1 and S2 polypeptides are colored in red and cyan with regions membrane-proximal from hinge 2 colored in white. The transmembrane domain of all chains is depicted in yellow. To design a Spike-Ferritin molecule, the C-terminal heptad repeat (residues 1140 to 1161) between Hinge 1 and 2 were aligned to an ideal heptad repeat sequence. Residues in the native S sequence which break this pattern are highlighted in red. These residues are also labeled and highlighted in red on the 3D-structure. Two engineered designs (1B-05 and 1B-06) are shown, with S end residue used to link to Ferritin, and heptad-repeat mutations colored green. (B) Schematic and 3D model of Spike Ferritin nanoparticle (SpFN). Differences between the native S sequence and the engineered nanoparticle are indicated on the schematic. A 3D-model of SpFN displaying eight trimeric Spikes was created using PDB ID 6VXX and 3EGM with the ferritin molecule shown in alternating grey and white. The nanoparticle is depicted along the 4-fold and the 3-fold symmetry axes of the ferritin. (C) RBD–Ferritin nanoparticle design and optimization. The RBD of SARS-CoV-2 (PDB ID:6MOJ) is shown in surface representation, with the ACE2 binding site outlined in dashed lines. Three hydrophobic regions of the RBD which were mutated for nanoparticle immunogen design are shown in light green surface, with residues in stick representation. The ACE2 binding site contains two of these regions, while a third hydrophobic patch near the C-terminus of the RBD is typically buried by S2 and part of S1 in the context of the trimer molecule. (D) Schematic and 3D model of an RBD–Ferritin nanoparticle. A modeled 24-mer nanoparticle displaying the RBD domain is depicted at the 3-fold symmetry axis of ferritin and colored green. Truncation points, linkers, and alterations made to the RBD sequence are indicated on the schematic. (E) Schematic and 3D model of an RBD–NTD–Ferritin nanoparticle. A modeled nanoparticle displaying RBD and NTD epitopes is depicted and colored according to the schematic. Truncation points, linkers, and alterations made to the native S sequence are indicated on the primary structure. (F) S1-Ferritin immunogen design. The SARS-CoV-2 S1 forms a hydrophobic collar around the N-terminal β-sheet of S2 (residues 689-676). S1-ferritin immunogen design required inclusion of this short stretch of S2 (colored cyan) attached by a linker. Terminal residues of the structured portions of S1 and S2 are labeled. (G) Schematic and 3D model of an S1–Ferritin nanoparticle. A modeled nanoparticle displaying RBD and NTD domains is depicted and colored according to the S1–ferritin schematic with truncation points and domain linkers indicated. See also Figure S1 and Table S1.

The first design category, Spike ferritin nanoparticle designs were based on a modified S with stabilizing prolines (K986P, V987P), removal of the furin cleavage site (RRAS to GSAS), and optimization of the coiled coil region between hinge 1 and hinge 2 of the ectodomain stalk (Turonova et al., 2020) to stabilize trimer formation on the Ferritin scaffold (Figure 1A and S1). The designs focused on (i) modification of the end of the S molecule (1137, 1208, 1154, or 1158), (ii) optimization of the coiled-coil region through extensions or repeats, (iii) removal of the coiled-coil region, (iv) removal of glycan 1158, (v) addition of heterogenous trimerization domains (GCN4, or foldon), or (vi) signal peptide sequence (Figure 1B, and Table S1).

The second design category, RBD ferritin nanoparticle designs used the SARS-CoV-2 RBD (residues 331-527) (Figure 1C) connected to the bullfrog-H. pylori chimeric ferritin (Kanekiyo et al., 2015) by a 6 amino acid linker (Figure 1D). The SARS-CoV-2 RBD contains a set of hydrophobic patches, including the ACE2 binding site, and a region located about residue 518 that is covered in the context of the intact S molecule. These regions were iteratively mutated to reduce hydrophobicity, and increase stability of the RFN molecules, expression levels, and antigenicity and immunogenicity.

The third design category, RBD-NTD ferritin molecules were based on addition of optimized RBD molecules in series with an NTD-Ferritin construct (residues 12-303) linked to the bullfrog-H. pylori chimeric ferritin molecule. The in-series, but reversed RBD-NTD design ensured distal displacement of the RBD molecule from the ferritin molecule (Figure 1E), promoting immune recognition of the RBD molecule with potential benefits for the production and stability of the nanoparticle.

The fourth design category, S1 ferritin design SARS-CoV-2 S (residues 12 - 676) (Figure 1F, and Table S1) was initially designed based on the MERS S1 immunogen which elicited protective immune responses (Wang et al., 2015). Subsequent designs focused on inclusion of a short region of SARS-CoV-2 S2 (residues 689-696) either using the connecting region that overlaps with the furin site, or by use of a short glycine-rich linker sequence (Figure 1F) to enable formation of the S1-Ferritin nanoparticle (Figure 1G).

### Characterization of SARS-CoV-2 S-domain ferritin nanoparticles

Ten S-ferritin constructs (Table S1) were initially designed and tested for expression, yield, nanoparticle formation, and antigenicity. S-ferritin nanoparticles were expressed in Expi293F cells for 3-5 days at 34 °C and 37 °C and purified by GNA lectin affinity chromatography. A subset of these constructs showed reasonable expression levels ranging from 0.5 to 5 mg/L media supernatant (Figure S3). pCoV1B-05 and pCoV1B-06-PL (SpFN) typically yielded over 5 mg/L with expression incubation set at 34 °C. Samples were assessed by SDS-PAGE, size-exclusion chromatography (SEC), dynamic light scattering (DLS), and negative-stain electron microscopy (neg-EM) to ensure intact protein was produced, and to assess the nanoparticle formation, and S morphology (Figure 2 and . For all SpFN constructs that showed expression as visualized by SDS-PAGE (Figure 2A), nanoparticles were observed by SEC, DLS, and neg-EM (Figure 2E, and 2F). In the case of SpFN and SpFN_1B-08 (Figure S2), the globular shape of the protruding S was clearly visible in both the TEM images and the 2D averages. In the case of the pCoV1B-05, the protruding S showed more of an “open” form in both the TEM images and the 2D averages.

**Figure 2.**
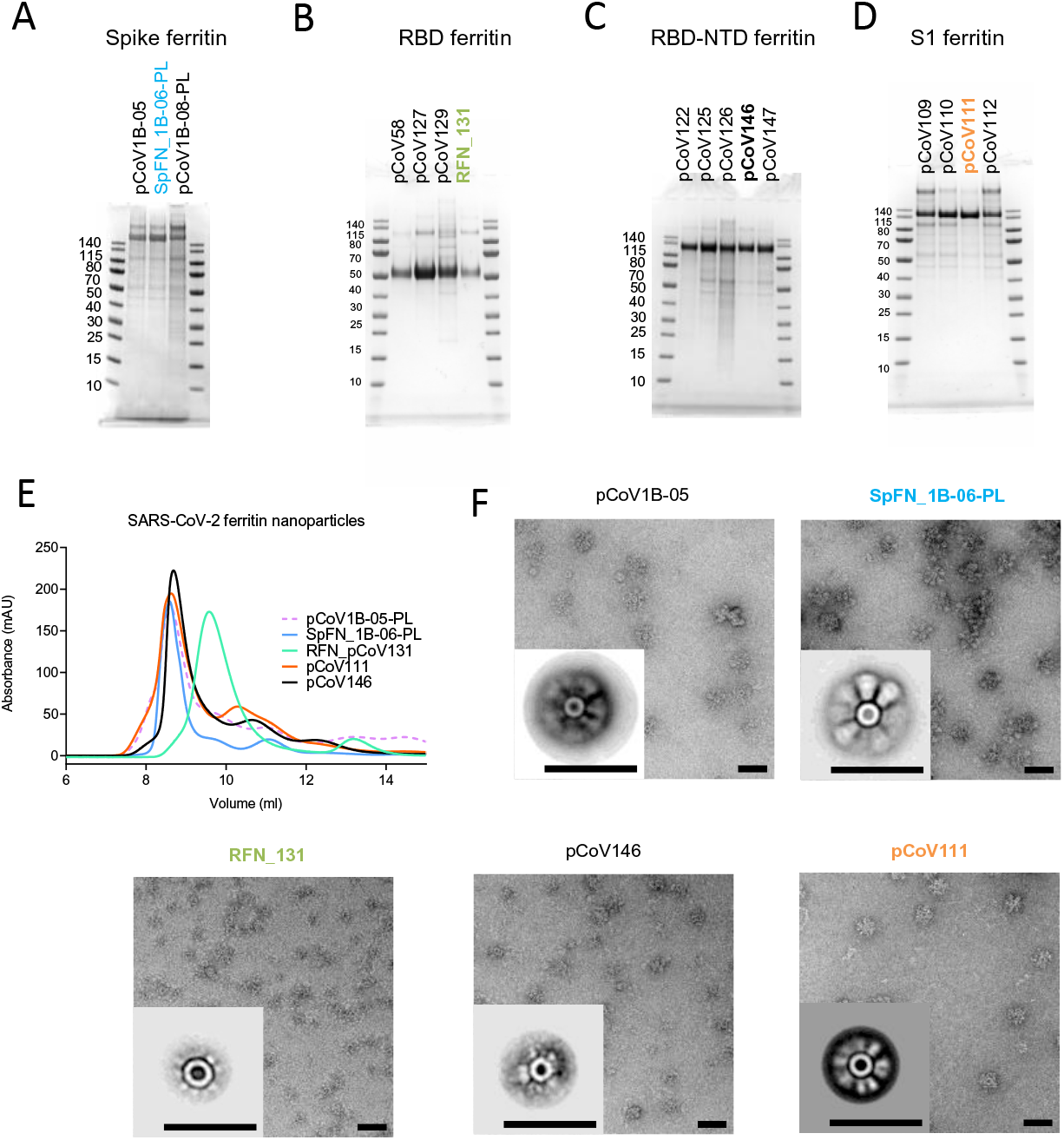
Antigenic and biophysical characterization of SARS-CoV-2 Spike-based ferritin nanoparticle vaccine candidates. SDS-PAGE of (A) Spike-Ferritin nanoparticle designs, (B) Receptor-binding domain-Ferritin nanoparticle designs, (C) S1-ferritin nanoparticle, and (D) RBD-NTD-Ferritin nanoparticles. Molecular weight standards are indicated in kDa. (E) Size-exclusion chromatography on a Superdex S200 10/300 column of representative SARS-CoV-2 S-based ferritin nanoparticles. (F) Negative-stain electron microscopy 2D class averages of purified nanoparticles. The black bars represent 50 nm. See also Figure S2 and S3.

Nanoparticles were assessed for nanoparticle formation, and assessed for antigenicity using biolayer interferometry against a set of poorly neutralizing (CR3022, SR1), and potently neutralizing (SR2, SR3, SR4, SR5) RBD-targeting antibodies. The different S-ferritin designs showed variable binding to the antibodies, with SpFN having the highest binding (Figure 3A and Figure S1).

**Figure 3.**
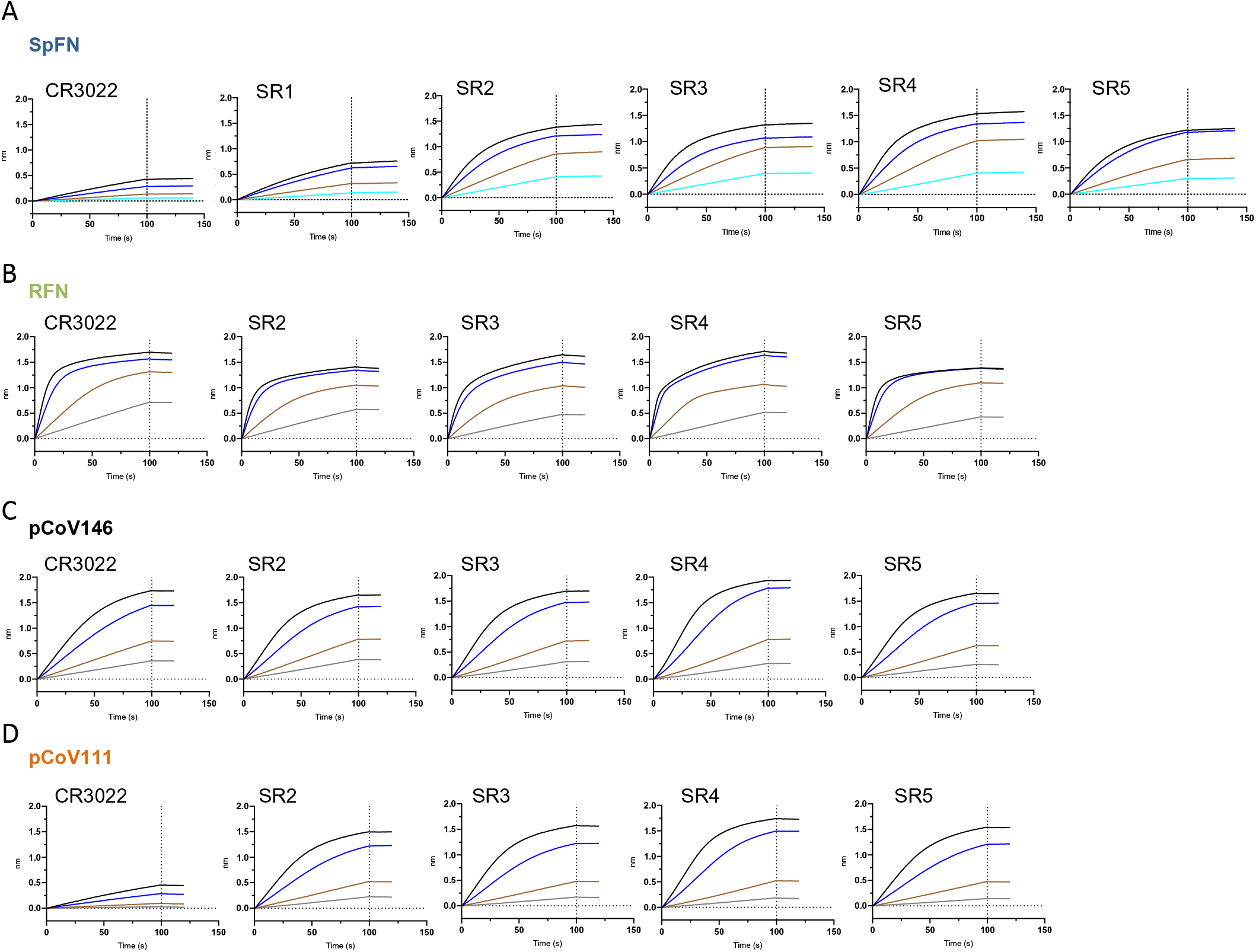
Antigenic characterization of select SARS-CoV-2 Spike-based ferritin nanoparticle vaccine candidates. Binding response of SARS-CoV-2 neutralizing antibodies to each of the lead candidates from the four design categories measured by biolayer interferometry with two-fold serial dilution of each antibody starting at 30 μg/ml). (A) Spike-Ferritin nanoparticle SpFN_1B-06-PL. (B) RBD-Ferritin pCoV131. (C) RBD-NTD-Ferritin nanoparticle pCoV146. (D) S1-Ferritin nanoparticle pCoV111. See also Figure S3.

Initial test expression of RBD-Ferritin constructs at 37 °C using either 293F or Expi293F cells showed low levels of expression. Reducing the cell expression temperature to between 30 °C to 34 °C following transfection rescued expression and enabled levels > 20 mg/L to be purified by NiNTA purification (Figure S3). However, analysis of the constructs by SEC, DLS, and neg-EM indicated that initial RBD-Ferritin constructs did not form fully intact nanoparticles (Figure S2). We hypothesized that designed variants with reduced RBD surface hydrophobicity would allow for improved nanoparticle yield. Screening through a set of variants using SDS-PAGE and SEC as primary indicators of nanoparticle formation allowed identification of constructs that readily formed nanoparticles (Figure 2 and Figure S1 and S2B). These molecules were also visualized by neg-EM, and showed clear formation of nanoparticles, with the protruding RBD domain visible on their surface in both TEM images and 2D class averages (Figure 2F). However, these constructs had a propensity to form soluble or insoluble aggregates and dramatically affected the ability to concentrate the samples. Addition of 5% glycerol to the NiNTA purified material, prior to SEC or other concentration steps, mitigated the aggregation issue and increased the nanoparticle formation as judged by SEC, and was confirmed by neg-EM. The RBD-Ferritin constructs showed very strong binding to the set of RBD-specific antibodies in all cases (Figure S3E). The level of binding was approximately twice that seen for the S-ferritin constructs, indicative of the exposed and accessible nature of the RBD epitopes (Figure 2G).

Due to the initial difficulty with S1-Ferritin nanoparticle constructs, we developed a set of engineered S1 constructs by artificially connecting the RBD domain by a short linker to NTD linked to the Ferritin molecule and denoted as RBD-NTD ferritin molecules. This multi-domain strategy resulted in good expression and nanoparticle formation. Using the information gained from the RBD surface optimization, we designed multiple constructs with variations in the RBD molecule to reduce surface hydrophobicity (Figure S1).

Antigenic analysis of these constructs showed that pCoV146 displayed robust antibody-binding (Figure 2F). The initial S1-Ferritin construct, pCoV68 (residues 12-676) yielded very low protein expression levels, even with reduced expression temperatures(Figure S3A). However, using the structure of the S-2P molecule (Wrapp et al., 2020; Walls et al., 2020), it was clear that a short segment of the S2 formed significant interactions with the S1 domain. Addition of this short region either using the natural sequence (with furin site removed) as in construct pCoV109, or by linking residues 689-696 with glycine-rich linkers as in construct pCoV111 allowed ∼ 1 mg/L of protein to be purified. Analysis of these constructs by SDS-PAGE, SEC, and neg-EM showed clear formation of the designed nanoparticles, and antigenic characterization showed binding of antibodies to the nanoparticle (Figure 2G).

Further structural analysis of the nanoparticle immunogens from each of the four design categories was carried out by determining 3D reconstructions from negative-stain electron micrographs (Figure 4). For each nanoparticle, a central sphere of approximately 12 nm corresponding to ferritin was resolved. S-domain antigens were located a short distance away from the central sphere and linker regions were unresolved likely due to their small size and flexibility. The SpFN_1B-06-PL reconstruction showed the stabilized S protruding from the ferritin molecule with a total diameter of approximately 44 nm (Figure 4A). The large size and distinct low-resolution features of the S ectodomain allowed for docking of a closed S-2P trimer model (PDB ID: 6VXX) into the trimer density, confirming the S was in the prefusion conformation. The ferritin-distal region of the S density was slightly weaker and likely reflects the heterogeneity in RBD-up conformations or slight openings of trimer visible in raw micrographs. Additionally, although the coiled-coil was unresolved, the distances between the density for S and ferritin matched the modeled coiled-coil length. Reconstruction of the three-dimensional RFN_131 EM map revealed two globular densities per asymmetric unit, suggesting that the RBD molecule was highly flexible on the surface of the ferritin sphere (Figure 4B). Similarly, the map of the RBD-NTD-ferritin fusion, pCoV146, showed two layers of globular densities, with a ferritin-proximal layer corresponding to the NTD domain and a more disordered layer for the RBD domain (Figure 4C). This particle was approximately 9nm larger in 2D and 3D than the single domain RFN molecule. The reconstruction of the S1-Ferritin fusion pCoV111 revealed a surprisingly ordered S1 density compared to the flexible RBD-NTD fusion, perhaps due to geometric constraints on the surface of the ferritin particle (Figure 4D). A density similar in shape to the S1 domain in the closed S2P trimer was resolved although it was slightly more compact, likely due to both overall flexibility of the S1 on the ferritin surface and RBD flexibility.

**Figure 4.**
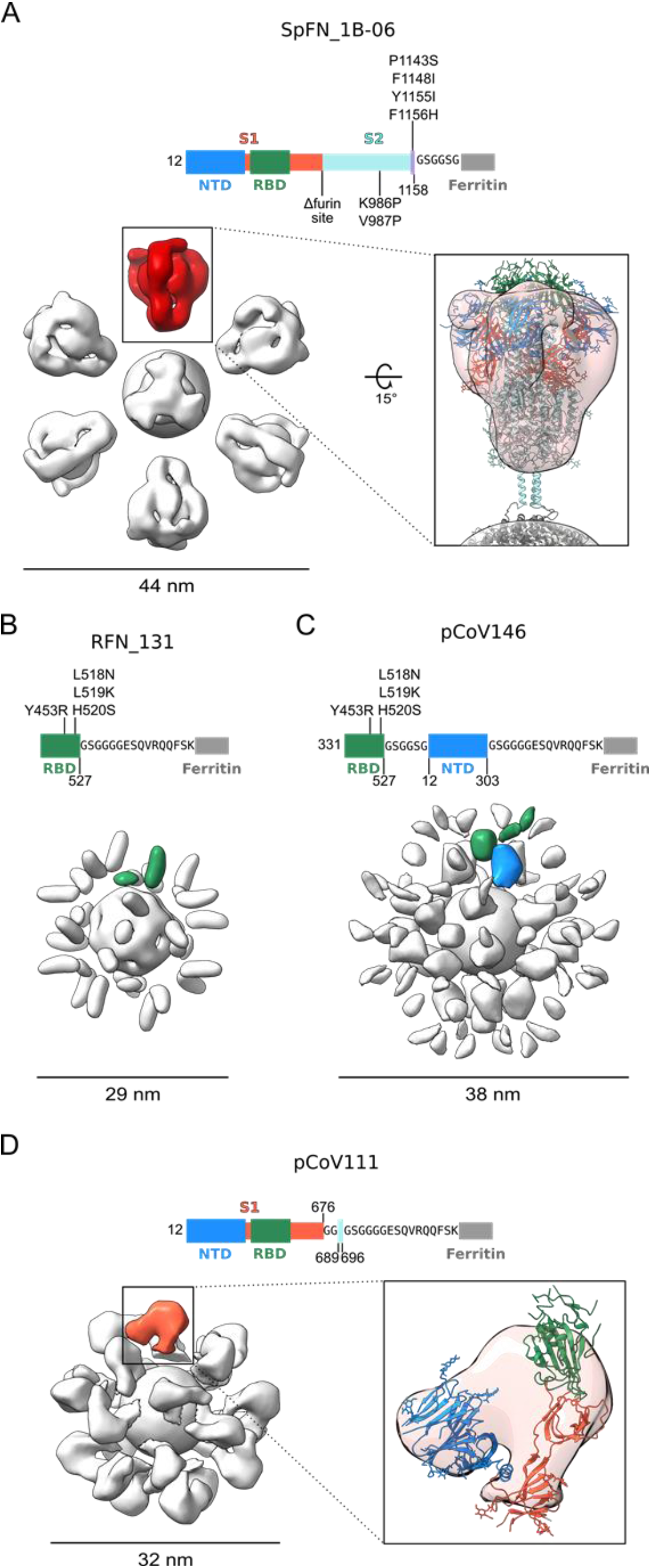
Negative-Stain Electron Microscopy 3D Reconstructions of SARS-CoV-2 Spike Domain-Ferritin Nanoparticles. Modifications made to native sequence and linkers used for each construct are shown in schematic diagrams. (A) Negative-stain 3D reconstructions with applied octahedral symmetry are shown with an asymmetric unit of non-ferritin density colored and the size of each particle indicated in nanometers. Spike trimer density, is colored in red, and a model of a SpFN trimer based on PDB 6VXX is shown docked into the negative-stain map and colored according to the sequence diagram. (B) Two non-ferritin densities per asymmetric unit were observed for RFN and are highlighted in green. These densities putatively correspond to the receptor-binding domain (RBD) but lack low resolution distinguishing features due to the small, globular shape of these domains. The presence of two densities is likely due to flexibility in the linker and heterogeneity in the RBD pose. (C) Two layers of densities were distinguishable for pCoV146, with the putative N-terminal domain (NTD) density of an asymmetric unit colored blue, proximal to the ferritin and two smaller, more flexible densities corresponding to RBD distal to the ferritin and colored green. (D) An asymmetric unit of non-ferritin density for pCoV111 is colored in orange and a monomer of S1 in the closed trimer state from PDB 6VXX is shown docked into the density with domains colored as in the sequence diagram. See also Figure S2 and Table S2.

### Immunogenicity of the four categories of SARS-CoV-2 S-domain ferritin nanoparticles in mice

To evaluate the immunogenicity of the SARS-CoV-2 ferritin-nanoparticles, we typically utilized two strains of mice (C57BL/6, and BALB/c), two adjuvants (ALFQ, and Alhydrogel^®^), and immunized mice three-times intramuscularly at 3-week intervals using a 10 μg dose. In total, we assessed 14 immunogens, two Spike ferritin immunogens, seven RBD ferritin immunogens, one S1-Ferritin construct, and four RBD-NTD ferritin immunogens (Table S2). Assessment in immunogenicity studies was based on iterative knowledge of immunogen physical and biochemical characteristics (Figure 2 and 3, and Figure S3), in conjunction with immunogenicity results from first-generation immunogens (Figure S1). This facilitated down-selection of lead immunogen candidates. Alhydrogel^®^ and ALFQ adjuvants were selected due to their history in human vaccine trials, safety profile, and previous performance alongside nanoparticle vaccine immunogens (NCT04296279). Alhydrogel^®^ contains aluminum hydroxide gel, while ALFQ is a liposome-based adjuvant containing the saponin QS-21, and synthetic Monophosphoryl Lipid A (3D-PHAD^®^). We assessed SARS-CoV-2 RBD- and S-binding, RBD-ACE2-inhibition, pseudovirus neutralizing antibody responses and authentic SARS-CoV-2 virus neutralization (Figure 4).

All four categories of immunogens elicited robust SARS-CoV-2 immune responses. In all cases tested, ALFQ was superior to Alhydrogel^®^ as an adjuvant for elicitation of binding and neutralizing responses (Figure S4). In addition, Alhydrogel^®^ led to a skewed antibody isotype immune response that was TH2 in nature, as opposed to the balanced immune response seen with ALFQ adjuvanted animals (Figure S4G). Immune responses seen in C57BL/6 mice were greater than for BALB/c mice after a single immunization, while binding and neutralizing antibody titers were comparable after a second or third immunization (Figure 5). In all cases, the third immunization did not dramatically increase the antibody levels induced by the S-domain ferritin nanoparticles.

**Figure 5.**
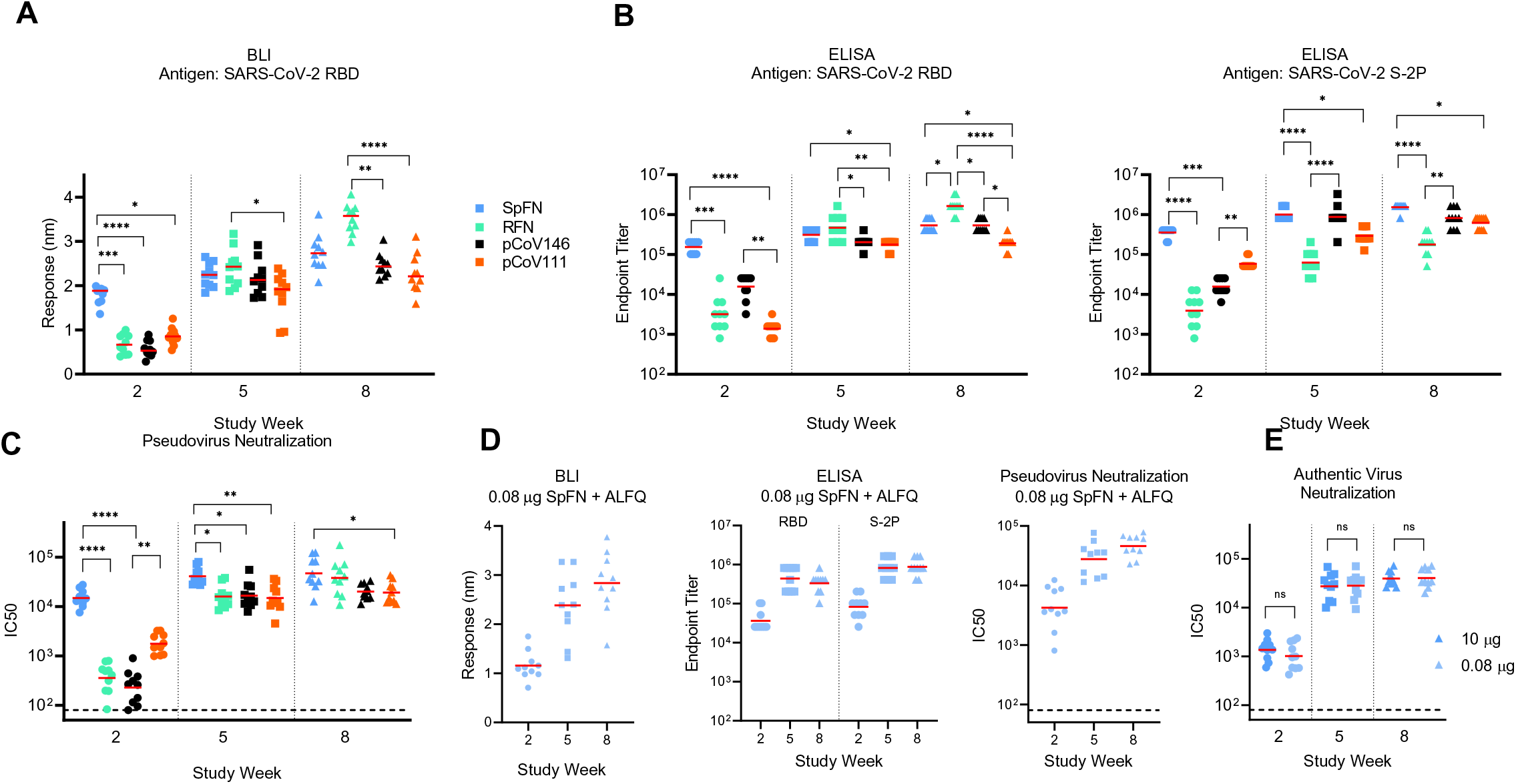
SARS-CoV-2 Spike-domain nanoparticle vaccine candidates elicit robust binding and neutralizing antibody responses in mice. (A) Biolayer interferometry binding of mouse sera to SARS-COV-2 RBD. Study week is indicated on the base of the graph. Mean value is indicated by a horizontal line. Statistical comparison at each timepoint was carried out using a a Kruskal-Wallis test followed by a Dunn’s post-test. (B) ELISA binding of mouse sera to SARS-COV-2 S-2P or RBD. Study week is indicated on the base of the graph. Geometric mean value is indicated by a horizontal line. Statistical comparison at each timepoint was carried out using a a Kruskal-Wallis test followed by a Dunn’s post-test. (C) SARS-COV-2 pseudovirus neutralization ID50 and ID80 values. Geometric mean value is indicated by a horizontal line. Statistical comparisons at each given timepoint was carried out using a Kruskal-Wallis test followed by a Dunn’s post-test. (D) Binding and pseudovirus neutralization of sera from mice immunized with 0.08 μg SpFN + ALFQ. (E) Authentic SARS-CoV-2 virus strain 2019-nCoV/USA_WA1/2020 neutralization ID50 and ID80 are shown for mice immunized with 10 or 0.08 μg SpFN + ALFQ. Geometric mean value is indicated by a horizontal line. Comparisons between dose group at each time point were carried out using a Mann-Whitney unpaired two-tailed non-parametric test, n=10 mice/group. In panels A – C, all immunogen groups at a given study timepoint were compared to each other. Only groups with statistically significant differences are indicated by a bar; all other groups did not show statistically significant differences. P values <0.0001 (****), <0.001 (***), <0.01 (**), or <0.05 (*). See also Figure S4 and S5, and Table S3 and S4.

The SpFN immunogen elicited a rapid RBD-binding, and pseudovirus neutralizing antibody response with ID50 geometric mean titer (GMT) >10,000 in C57BL/6 mice and >1,000 after a single immunization (Figure 5C and Figure S4). This rapid neutralizing immune response after one immunization was significantly higher than seen with RBD ferritin or RBD-NTD ferritin immunogens. Following a second immunization, both strains of SpFN-vaccinated mice showed ID50 GMT >10,000 and ID80 GMT >5,000 for both mouse types.

RBD ferritin immunogens elicited robust RBD and S-binding responses, ACE2-inhibition, and pseudovirus neutralization ID50 GMT >10,000 in both mice strains after two immunizations (Figure 5A-C). We assessed seven RFN immunogen designs in immunogenicity studies (Table S3) and selection for animal immunogenicity experiments was largely based on nanoparticle stability, expression levels, aggregation, and antigenic profile (Figure 2-3 and Figure S2, S3). Based on these criteria, pCoV131 (RFN) was extensively assessed, and after three immunizations, showed pseudovirus neutralization responses that were comparable or exceeded that seen for the SpFN_1B-06-PL immunogen. Of note, the RFN immunogens elicited substantial S binding responses that were highly comparable to that of other immunogens that contained additional S domains (Figure 5B).

In a pattern similar to that seen for the RBD ferritin immunogens, both the RBD-NTD ferritin and the S1 ferritin immunogens elicited binding responses, and detectable pseudovirus neutralization after a single immunization that were increased by the second immunization to give ID50 GMT values >10,000, and ID80 GMT titers ∼5,000 (Figure 5A-C and Figure S4).

Given the rapid elicitation of immune responses after a single immunization by SpFN_1B-06-PL, we further characterized this immunogen in a dose-ranging study (Figure S5 and Figure 5D and 5E). In five-fold dilution steps, we reduced the SpFN_1B-06-PL immunogen from a 10 μg dose down to a 0.0032 μg dose (3,125-fold reduction) with the full ALFQ adjuvant dose. Antibody binding responses were assessed for the full dose range by ELISA to S and RBD, and binding responses were elicited at all dose concentrations tested. We then further assessed the 0.08 μg dose (125-fold reduction from the 10 μg dose) with all our immunogenicity assays (Figure 5D and Figure S5G). At this dose, the immune response was comparable to that seen for the typical 10 μg dose. In addition, we assessed both the 10 and 0.08 μg SpFN_1B-06-PL vaccinated-mouse serum for authentic SARS-CoV-2 live virus neutralization. At both doses, in both mouse strains, a single immunization elicited ID50 GMT of ∼ 1,000, while the second immunization boosted this response more than ten-fold. The subsequent third immunization showed a modest boost effect. At each of these study time points, there was no difference between the doses in C57BL/6 mice, while differences in BALB/c ID80 GMT were seen at week 2 (higher for the 10 μg dose) and week 8 (higher for the 0.08 μg dose). Overall, these studies demonstrated robust immunogenicity of four categories of SARS-CoV-2 S-domain ferritin nanoparticles.

### Protective Immunity in K18-hACE2 transgenic mice against SARS-CoV-2 Challenge

Given the neutralizing antibody response seen with SpFN, and RFN, and the different design of these two immunogens, we chose to assess antibodies from these animals in a lethal SARS-CoV-2 challenge model using K18-hACE2 transgenic mice. The dose of SARS-CoV-2 virus was titrated to establish significant weight loss and pathology following infection with WA strain (Figure S6). Given previously described studies (Zheng et al., 2021), we sought to assess the vaccine-elicited antibody responses at levels starting at about 1,000 and we passively transferred three different amounts of purified IgG from either SpFN- or RFN-immunized mice 24 h prior to infection with SARS-CoV-2 (Figure 6A, 6B). The control groups included a PBS group and a group that was passively transferred with naïve mouse IgG.

**Figure 6.**
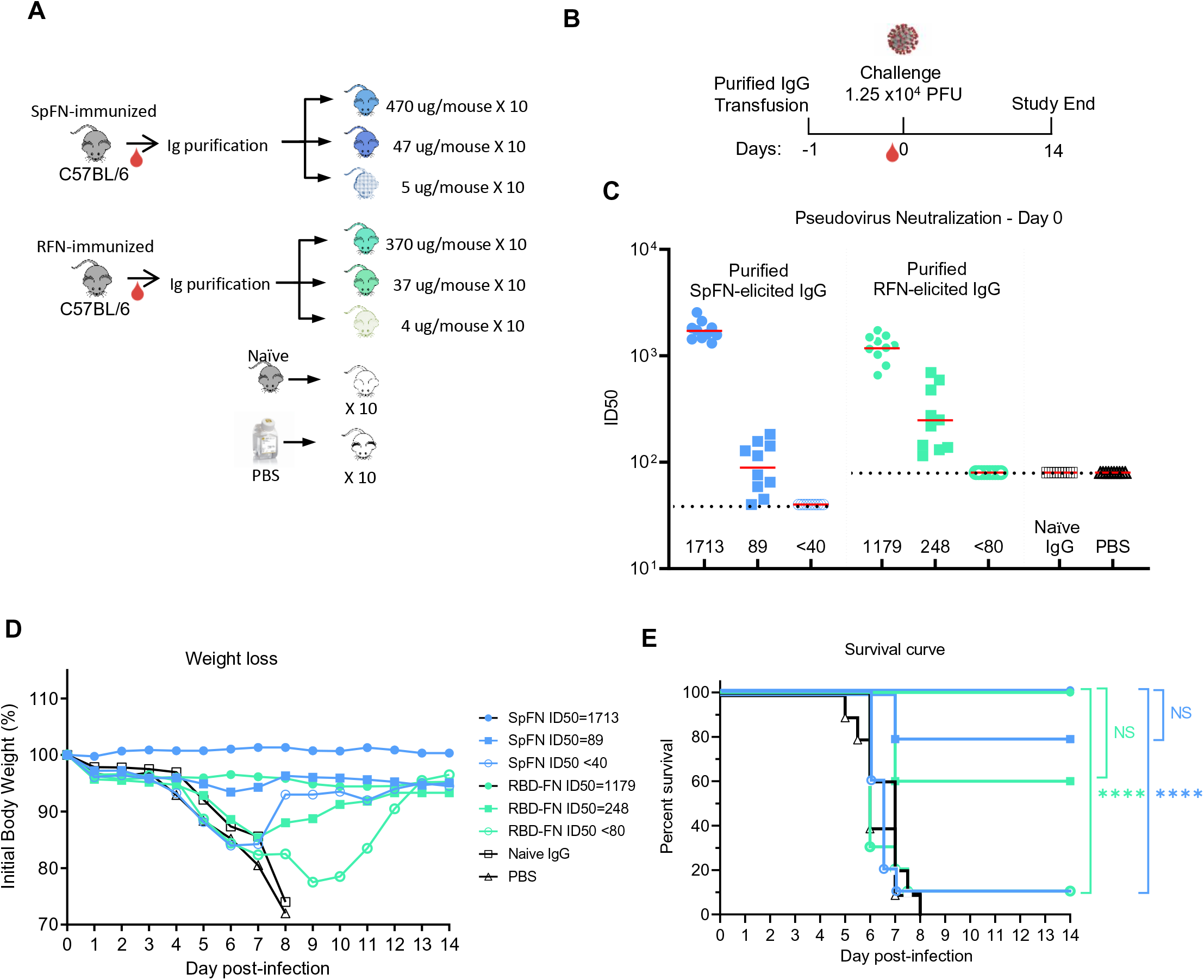
SpFN- and RBD-FN protective immunity in K18-hACE2 transgenic mice. (A) IgG was purified from SPFN- or RFN-vaccinated mouse sera and passively transferred at specific IgG amounts ranging from 4 - 470 μg/mouse in a final volume of 200 μl. Control naïve mouse IgG was formulated at 2 mg/ml. (n=10/group, 5 female, 5 male). (B) Mouse challenge study schematic. K18-hACE2 mice (n=10/group, 5 female, 5 male) received control IgG (black), PBS (gray), and purified IgG, one day prior to challenge with 1.25 × 10^4^ PFU of SARS-CoV-2. (C) SARS-CoV-2 pseudovirus neutralization ID50 titers of mouse sera at study day 0. (D) Percentage of initial weight of K18-hACE2 mice for the 8 study groups. Legend is shown in panel E. (E) Survival curves of K18-hACE2 mice with groups indicated based on animal vaccination group and the pseudovirus ID50 neutralization values. Statistical comparisons were carried out using Mantel-Cox test followed by Bonferroni correction. See also Figure S6.

Animal weight was measured twice daily for 14 days after challenge, and animals that lost > 25% weight during the study were euthanized. All animals that received the highest amount of antibody (470 μg SpFN-derived, or 370 μg RFN-derived) showed neutralization ID50 GMT titers of 1,713 and 1,179 respectively (Figure 6C). All animals in these two groups showed minimal weight loss (Figure 6D), and all animals survived the study (Figure 6E). In the two groups that received either 47 μg SpFN- or 37 μg RFN-derived antibody, neutralization ID50 GMT titers were 89, and 248 respectively. Even with these modest antibody transfer amounts and the relatively low neutralization titers, most animals were protected from weight loss and death. In the SpFN-47 μg group, only two animals showed severe weight loss, while in the RFN-37 μg group, most animals showed some weight loss during the first study week, but all recovered. In contrast, mice that received the lowest amount of purified IgG from SpFN- or RFN-vaccinated animals, did not show any neutralizing antibody titers at the day of infection. The mice in these two groups showed significant weight loss, and 9/10 animals in each group were euthanized by day 9 of the study. In a similar pattern, all animals from the naïve IgG and PBS groups suffered weight loss and were euthanized by study day 8. In summary, these data show that low amounts of passively transferred antibodies from SpFN or RFN vaccinated animals can protect mice from a lethal challenge with SARS-CoV-2.

### Vaccine-elicited broadly cross-reactive antibody responses against SARS-CoV-1 and SARS-CoV-2

SARS-CoV-2 variants that are more transmissible and appear to be more lethal continue to emerge even in the midst of rapid vaccine roll out and public-health measures. Given the robust binding, pseudovirus and authentic virus neutralization titers against the original SARS-CoV-2, that were elicited by the S-domain ferritin nanoparticle immunogens, we assessed the immunized mouse sera for binding and pseudovirus neutralization to the VoC (Figure 7). Using study week 10 sera from mice immunized with the four categories of immunogens SpFN_1B-06-PL, RFN_131, pCoV146, and pCoV111 (S1-Ferritin), we assessed binding to a panel of variant RBD molecules containing K417N, E484K, N501Y, and combinations of these mutations (Figure 7A). These mutations match to the RBD sequence seen in the B.1.351, B.1.1.7, and P.1 SARS-CoV-2 strains. In all cases, robust binding to the RBD molecules were observed, with minimal change in overall binding when compared to the original RBD molecule. RFN-immunized mouse sera showed reduced binding to the E484K, N501Y double mutant, but increased binding to the K417N variant. Analysis of the sera from SpFN-, RFN, or pCoV111-immunized mice for pseudovirus neutralization of VoCs B.1.1.7 and B.1.351 showed minimal changes in the neutralization levels, with ID50 values > 2,000 for all strains (Figure 7B).

**Figure 7.**
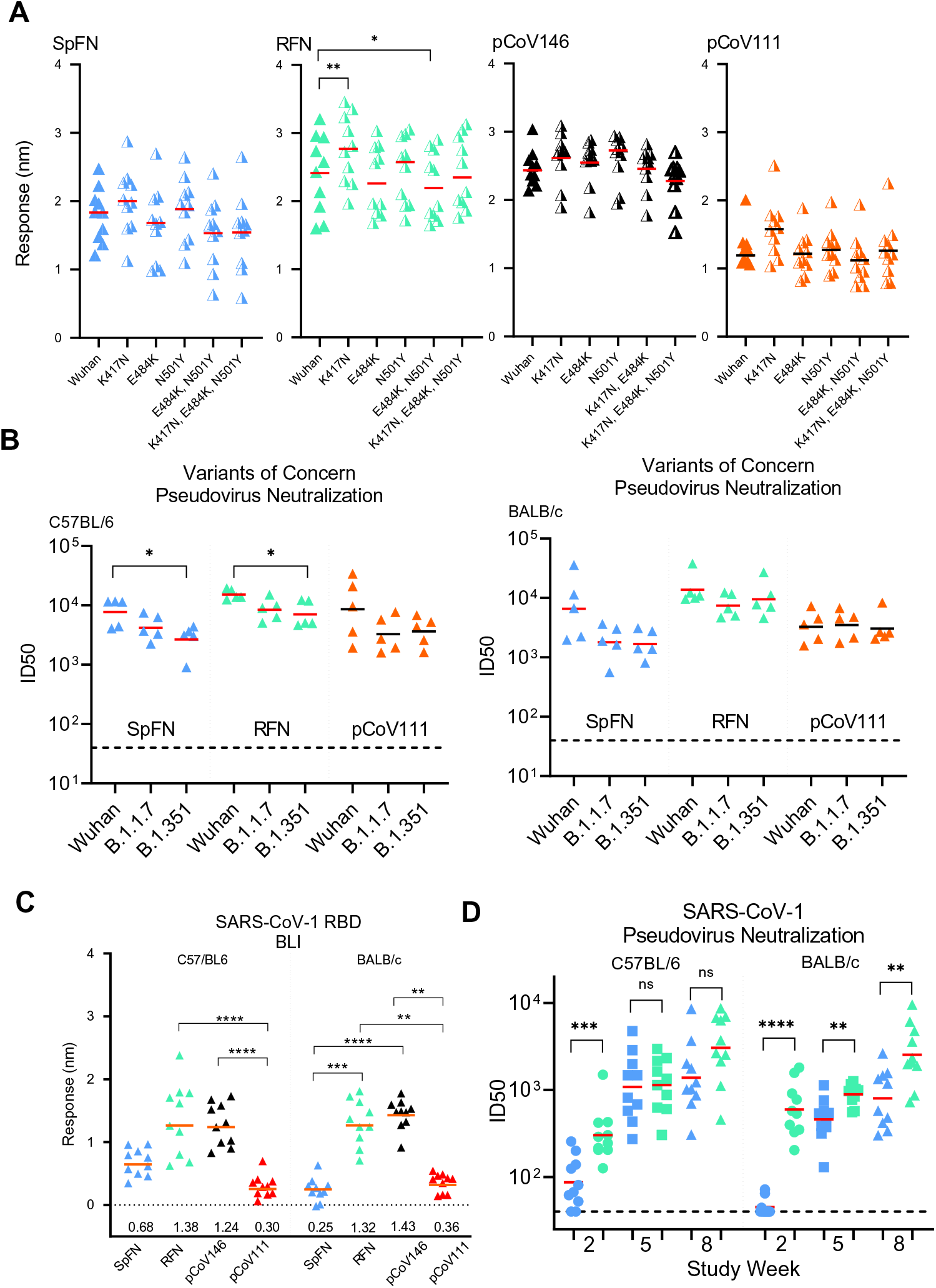
SARS-CoV-2 Spike-domain nanoparticle vaccine candidates elicit robust antibody binding responses and neutralizing activity against SARS-CoV-2 VoC and SARS-CoV-1. (A) Biolayer Interferometry binding of study week 10 immunized C57BL/6 mouse serum to SARS-CoV-2 RBD, and SARS-CoV-2 RBD variants. Immunogens are indicated at the top left of each graph. Mean values are indicated by a horizontal line, n=10, Significance was assessed using a Kruskal-Wallis test followed by a Dunn’s post-test. (B) Pseudovirus neutralization (ID50 values) of study week 10 immunized C57BL/6 and BALB/c mouse serum to SARS-CoV-2 Wuhan-1, B.1.1.7, and B.1.351 pseudotyped viruses. Immunogens are indicated at the base of each graph. Geometric mean values are indicated by a horizontal line, n=5, statistical significance for each immunogen was assessed using a Kruskal-Wallis test followed by a Dunn’s post-test. (C) Biolayer Interferometry binding of study week 10 immunized C57BL/6 and BALB/c mouse serum to SARS-CoV-1 RBD. Immunogens are indicated at the base of each graph. Mean values are indicated by a horizontal line, n=10, Significance was assessed using a Kruskal-Wallis test followed by a Dunn’s post-test. (D) Pseudovirus neutralization (ID50 values) of study week 10 immunized C57BL/6 and BALB/c mouse serum to SARS-CoV-1 Urbani strain pseudotyped viruses. Data related to SpFN and RFN are colored blue and green respectively. Statistical comparisons between SpFN and RFN responses at each time point were carried out using a Mann-Whitney unpaired two-tailed non-parametric test. Immunogens are indicated at the base of each graph. Geometric mean values are indicated by a horizontal line, n=10, P values <0.0001 (****), <0.01 (**) or <0.05 (*). See also Figure S4 and S5.

Analysis of the mouse sera for binding or neutralization of SARS-CoV-1 showed that RFN-immunized mice elicited the highest SARS-CoV-1 RBD binding response (Figure 7C). In addition, to RBD binding, we also observed SARS-CoV-1 ACE2-RBD inhibitory activity with the SpFN-immunized mice (Figure S5). We further assessed the SpFN- or RFN-immunized mouse sera for neutralization against SARS-CoV-1 using the pseudovirus assay (Figure 7D). We observed robust neutralization levels with ID50 > 1,000 for the SpFN_1B06-PL or the RFN_131 immunized animals. In general, the RFN molecule elicited higher SARS-CoV-1 neutralizing responses compared to the SpFN-immunized animals. Together, these data demonstrate that the S-domain ferritin nanoparticles elicit broadly neutralizing and cross-reactive antibody responses against VoC and heterologous SARS-CoV-1.

## DISCUSSION

Since the emergence of SARS-CoV-2 in late 2019, multiple vaccines have been developed that elicit robust and protective immune responses in small animals, non-human primates, and humans. This includes a set of mRNA-based vaccines (NIH-Moderna Pfizer-BioNTech), viral vector vaccines (J&J, Astra Zeneca), and a nanoparticle-like vaccine (Novavax) (Bangaru et al., 2020) that are starting to be distributed worldwide, and include SARS-CoV-2 S as the major vaccine component. In addition, next generation SARS-CoV-2 vaccine candidates are beginning to reveal strong immunological results in pre-clinical studies (Saunders et al., 2021; Walls et al., 2020). These protein-based nanoparticle platforms paired with powerful adjuvant systems provide multiple advantages in the ability to protect against emerging variants (Moyo-Gwete et al., 2021; Wibmer et al., 2021). Nanoparticle vaccines may be critical for specific high-risk professions, or in populations where immune response titers are of particular importance (Atyeo et al., 2021) including the elderly (Collier et al., 2021) or immunocompromised (Boyarsky et al., 2021). The utility of highly stable vaccines that can elicit high neutralizing antibody titers after a single immunization, or vaccines that can be easily re-purposed for specific populations or as boosting immunogens is likely to help the long-term strategy for global COVID-19 vaccination.

Trimer-functionalized ferritin vaccines have been effective at eliciting neutralizing antibodies against class I fusion proteins, including influenza haemagglutinin (Kanekiyo et al., 2013; Kelly et al., 2020) and HIV envelope (He et al., 2016; Sliepen et al., 2015) as well as engineered nanoparticles including RBD-12GS-I53-50 (Walls et al., 2020). For example, in the case of respiratory syncytial virus, a stabilized prefusion Fusion glycoprotein vaccine based on subtype A can naturally elicit potent neutralizing antibody and T cell responses against the heterologous subtype B in animals and humans (Crank et al., 2019; Joyce et al., 2019). Here we developed a set of SARS-CoV-2 S-domain ferritin nanoparticle vaccines using structure-based vaccine design that recreate the structural and antigenic profile of S. These immunogens elicit antibodies with potent S-binding activity, hACE2-blocking activity, and potent neutralization activity against the homologous virus. In all four immunogen-design groups, antibody responses target the RBD domain and this response significantly contributes to the high neutralization responses. Additionally, dose-sparing immunization experiments show that significant antigen reduction can still elicit potent antibody responses, while simultaneously also showing robust levels of neutralizing antibodies against the heterologous SARS-CoV-1. This heterologous immune response is reminiscent of broad immune responses seen with Ferritin-HA immunogens (Kanekiyo et al., 2013), and demonstrate how nanoparticle immunogens can enhance the quality of the humoral immune response. Naturally occurring nanoparticle vaccines such as Yellow Fever 17D vaccine (Collins and Barrett, 2017) and human papillomavirus virus-like particle (VLP) vaccines (Lowy and Schiller, 2006) elicit robust and long-lived immune responses. For SARS-CoV-2 vaccine development, nanoparticles performed well in mouse (Keech et al., 2020; Walls et al., 2020) and non-human primate studies (Brouwer et al., 2021), with a designed S-Ferritin nanoparticle also resulting in robust immunogenicity in mouse studies (Powell et al., 2021).

Furthermore, the passive-transfer of vaccine-elicited purified antibody prevented death and significant weight loss in a high-dose SARS-CoV-2 challenge in the K18-hACE2 mouse model at neutralizing antibody levels that were routinely exceeded by SpFN- or RFN-vaccination. We also transferred very low amounts of antibody to the K18-hACE2 mice to assess for antibody dependent enhancement, and we saw no indication of faster weight loss, or enhanced symptoms in the mice. The fact that antibody amounts readily elicited after a single immunization are highly protective in this challenge model, and that low antibody levels do not enhance disease, highlights the promise of these vaccine candidates. A set of companion papers also maps out the cellular immune response and highlights the protective effect of SpFN and RFN in high-dose SARS-CoV-2 challenge studies (Joyce et al., 2021; King et al., 2021). Given the rapid elicitation of SARS-CoV-2 immune responses after a single immunization and the highly protective responses seen in the K18-hACE2 model, SpFN_1B-06-PL has been produced under current Good Manufacturing Practice (cGMP) conditions and is under assessment in an ongoing phase I clinical trial (NCT04784767).

The immune responses elicited by the ferritin nanoparticles with the adjuvant ALFQ were consistently superior to that seen with the aluminum hydroxidebased Alhydrogel adjuvant. This result is consistent with other studies indicating that aluminum hydroxide is sub-optimal at inducing SARS-COV-2 neutralizing antibody responses. The components of the ALFQ adjuvant including QS-21, are used in multiple industrial processes and scaled up for future advanced clinical trials. The COVID-19 pandemic has set many precedents in regard to vaccine development speed, novel platforms, and should garner a new age of vaccine development utilizing advanced antigens and adjuvants to train the immune response for increased protection.

Here, we utilized structure-based design to create four categories of immunogens using the ferritin-nanoparticle platform. Each of these different designs and the underlying development processes provide a greater understanding and framework for ongoing and future pan-coronavirus vaccine design and development. The design information outlined here can be readily transferred for emerging CoV pathogens or other ubiquitous “common-cold” coronaviruses. The utilization of the SARS-CoV-2 nanoparticle immunogen provided immunogenicity against variants of concern and the heterologous SARS-CoV-1 and has implications for vaccination efforts against putative zoonotic emergences.

## ACKNOWLEDGEMENTS

We thank Sandhya Vasan, Mihret Amare, Suzanne Mate, Paul Scott, and Sharon P. Daye for programmatic support and planning, Nathaniel Jackson for cell culture maintenance; Erin Kavusak, and Jonah Heller for support with performance of the neutralization assays. Research was conducted in compliance with the Animal Welfare Act and other federal statutes and regulations relating to animals and experiments involving animals and adheres to principles stated in the *Guide for the Care and Use of Laboratory Animals*, NRC Publication, 1996 edition. This work was funded by the US Defense Health Agency, the US Department of the Army, and a cooperative agreement between The Henry M. Jackson Foundation for the Advancement of Military Medicine, Inc., and the US Department of Defense (W81XWH-18-2-0040). This study also was supported by grants from NIH (R01 AI157155) J.B.C. is supported by a Helen Hay Whitney Foundation postdoctoral fellowship. Material has been reviewed by the Walter Reed Army Institute of Research. There is no objection to its presentation and/or publication. The views expressed are those of the authors and should not be construed to represent the positions of the U.S. Army or the Department of Defense.

## AUTHOR CONTRIBUTIONS

M.G.J. and K.M. designed the study. M.G.J., P.V.T., and K.M. designed the immunogens. M.G.J., W-H.C., M.C., R.S.S., A.H., P.V.T., R.E.C. W.C.C., C.E.P., E.J.M., E.M., A.A., C.S., J.B.C., Y.L., A.A., J.K., T.O., L.R., A.G., C.W., J.C., L.M.-R., C.K., N.G., Z.V., D.McC., Z.B., J.K., S.S., O.J., V.D., S.M., U.T., C.B.K., M.Z., H.K., W.W., M.A.C., D.K.D., L.W.K., T.J.L., S.E.M., S.J.K., S.M., V.R.P., W.W.R., N.deV., M.S.D., G.D.G., and M.Rao performed protein purification, biophysical assays, immunologic assays and animal studies. Z.B., M.Rao, G.R.M., and A.And. designed and provided the adjuvants. S.R., P.M.M. and M.T.E. provided the SR1-5 antibodies. M.G.J., W.H.C., R.S.S., A.H., P.V.T., R.E.C., C.S., A.Ahm., L.W., Z.B., W.W., W.W.R., M.Ro., N.deV., M.S.D., G.D.G., M.Rao, N.L.M. and K.M. analyzed and interpreted the data. M.G.J. wrote the paper with assistance from all coauthors.

## STAR METHODS

Detailed methods are provided in the online version of this paper and include the following:

KEY RESOURCES TABLE
RESOURCE AVAILABILITY

Lead contact
Materials availability
Data and code availability statement
METHOD DETAILS

Immunogen Modeling and Design
DNA plasmid construction and preparation
Protein expression and purification
Negative-stain Electron microscopy
Octet Biolayer Interferometry binding and ACE2 inhibition assays
Mouse immunization
Immunogen-Adjuvant preparations
Enzyme Linked Immunosorbent Assay
SARS-CoV-2 and SARS-CoV-1 pseudovirus neutralization assay
Authentic SARS-CoV-2 virus neutralization assay
K18-hACE2 transgenic mouse passive immunization and challenge
QUANTIFICATION AND STATISTICAL ANALYSIS

### RESOURCE AVAILABILITY

#### Lead contact

Further information and requests for resources and reagents should be directed to and will be fulfilled by the Lead Contact, M. Gordon Joyce.

#### Materials Availability

All reagents will be made available on request after completion of a Materials Transfer Agreement.

#### Data and Code Availability

All data supporting the findings of this study are found within the paper and its Supplementary Information and are available from the Lead Contact author upon request.

## List of supplementary figures

**Figure S1.**
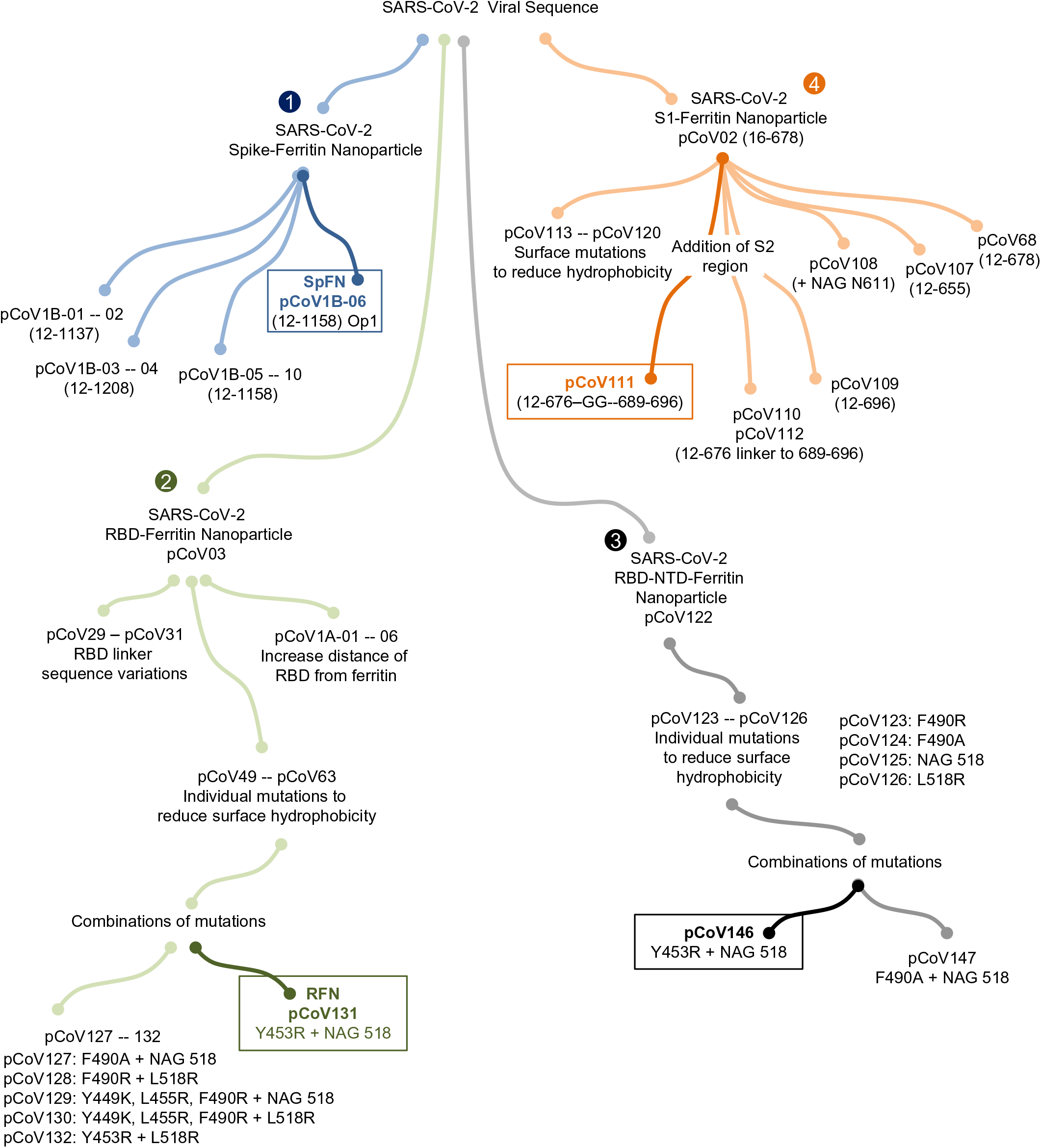
Structure-based design of SARS-CoV-2 Spike-based ferritin nanoparticle immunogens and design pipeline, related to Figure 1. Four ferritin nanoparticle immunogen designs were developed focused on (1) Spike ferritin nanoparticles (blue), (2) RBD ferritin nanoparticles (green), (3) RBD-NTD ferritin nanoparticles (black), and (4) S1 ferritin nanoparticles (orange). The design iterations and concepts are indicated, along with select mutations and design name. Lead vaccine candidates from each category are highlighted.

**Figure S2.**
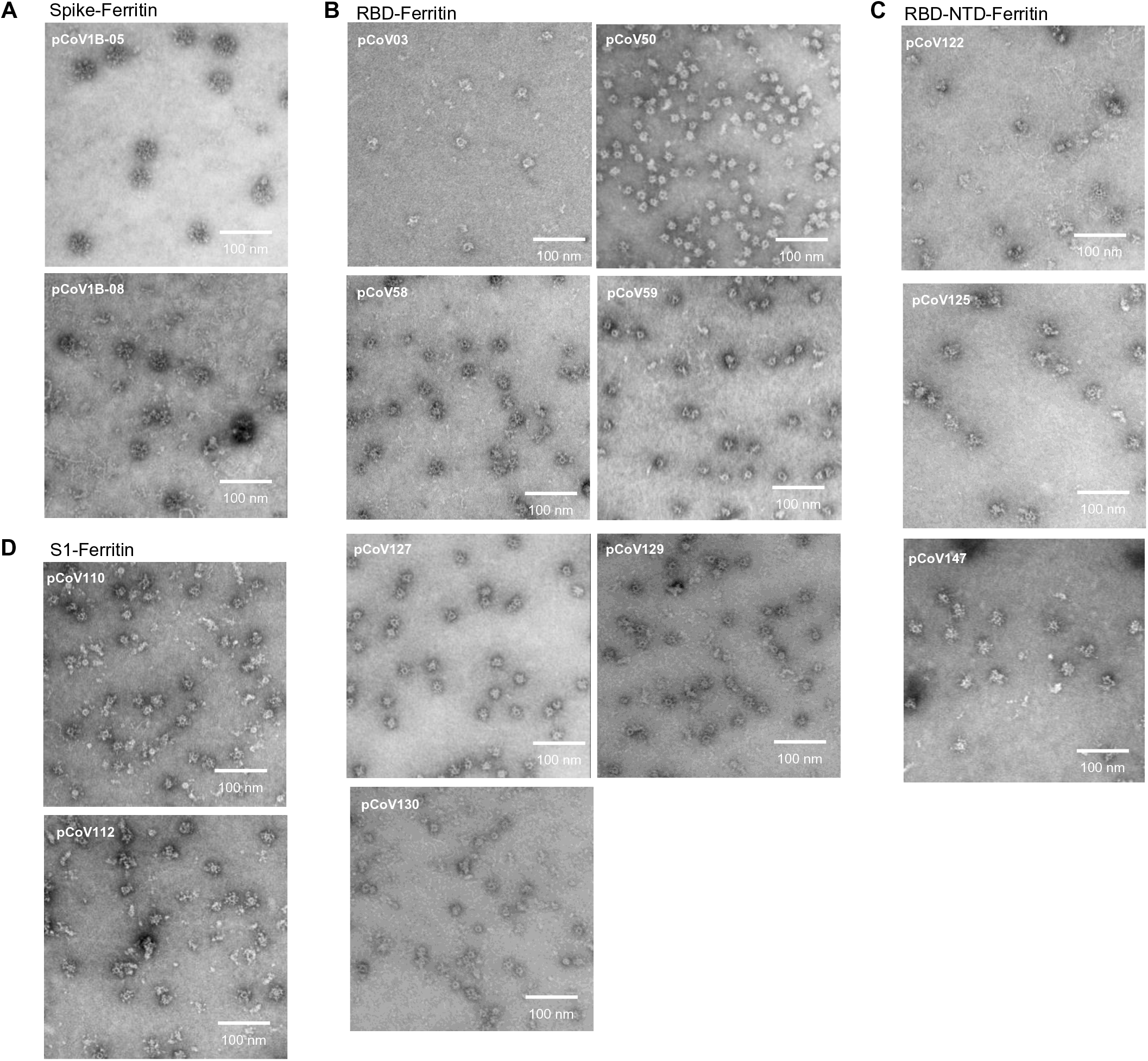
Negative-stain electron microscopy 2D micrographs of SARS-CoV-2 ferritin nanoparticle-based vaccine candidates, related to Figure 2 and 4. Negative-stain electron microscopy 2D micrographs. The white scale bars represent 100 nm. (A) Spike ferritin nanoparticles pCoV1B-05 and pCoV1B-08. (B) RBD ferritin nanoparticles pCoV03, pCoV50, pCOV58, pCoV59, pCoV127, pCoV129, pCoV130, pCoV131 (C) RBD-NTD ferritin nanoparticles pCoV122, pCoV125, pCoV147 (D) S1 ferritin nanoparticle pCoV110 and pCoV112.

**Figure S3.**
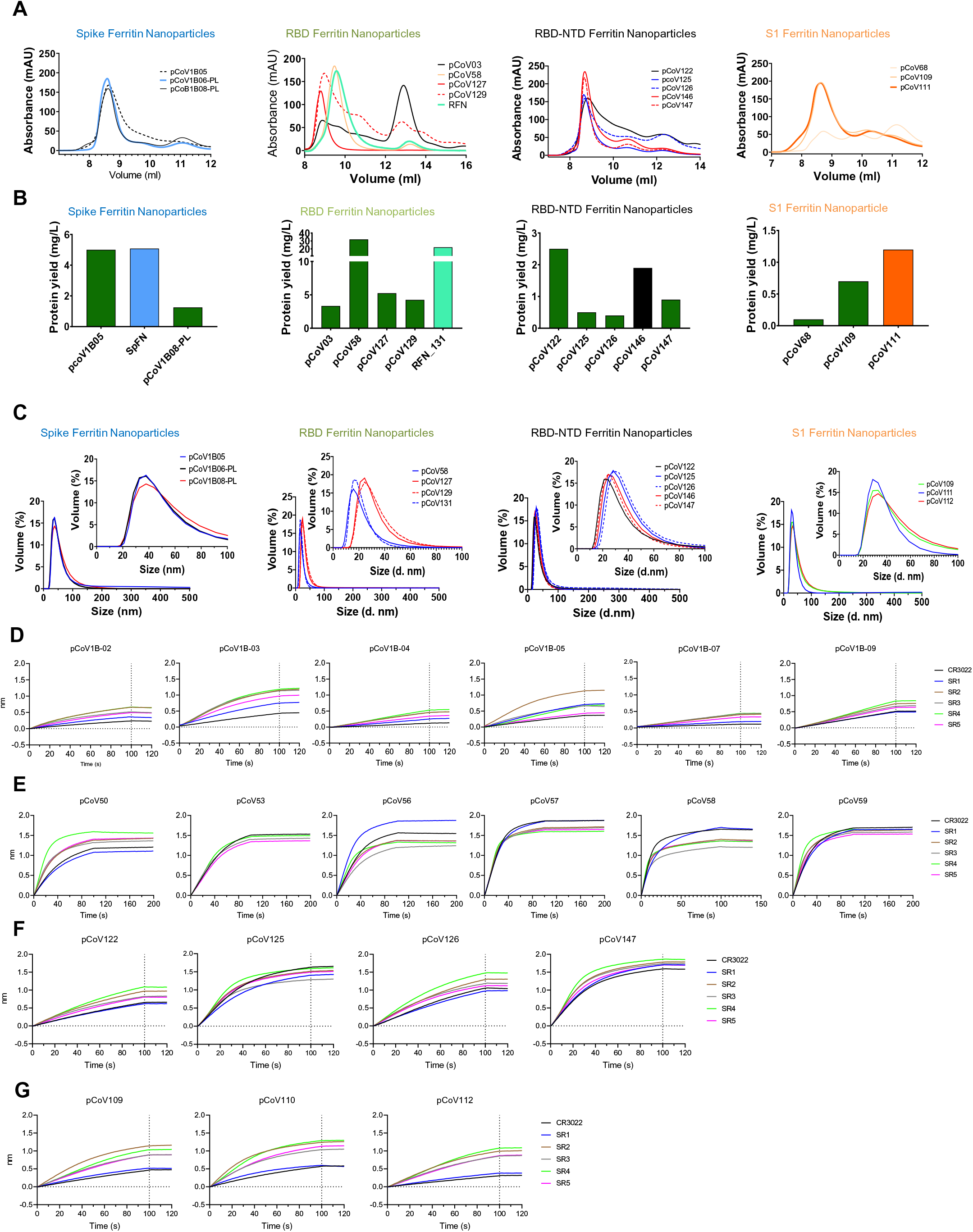
Biophysical and antigenic characterization of S-domain ferritin nanoparticle immunogens, related to Figure 2 and 3. (A) Size-exclusion chromatography on a Superdex S200 10/300 column of representative SARS-CoV-2 Spike-based ferritin nanoparticles from the four design categories. (B) Expression levels (mg/L supernatant) of representative SARS-CoV-2 Spike-based ferritin nanoparticles. (C) Dynamic light scattering analysis of representative SARS-CoV-2 Spike-based ferritin nanoparticles. (D) Spike ferritin nanoparticles (E) RBD ferritin, (F) RBD-NTD ferritin and (G) S1 ferritin nanoparticles were assessed for binding to a set of neutralizing antibodies (concentration = 30 μg/ml) by biolayer interferometry.

**Figure S4.**
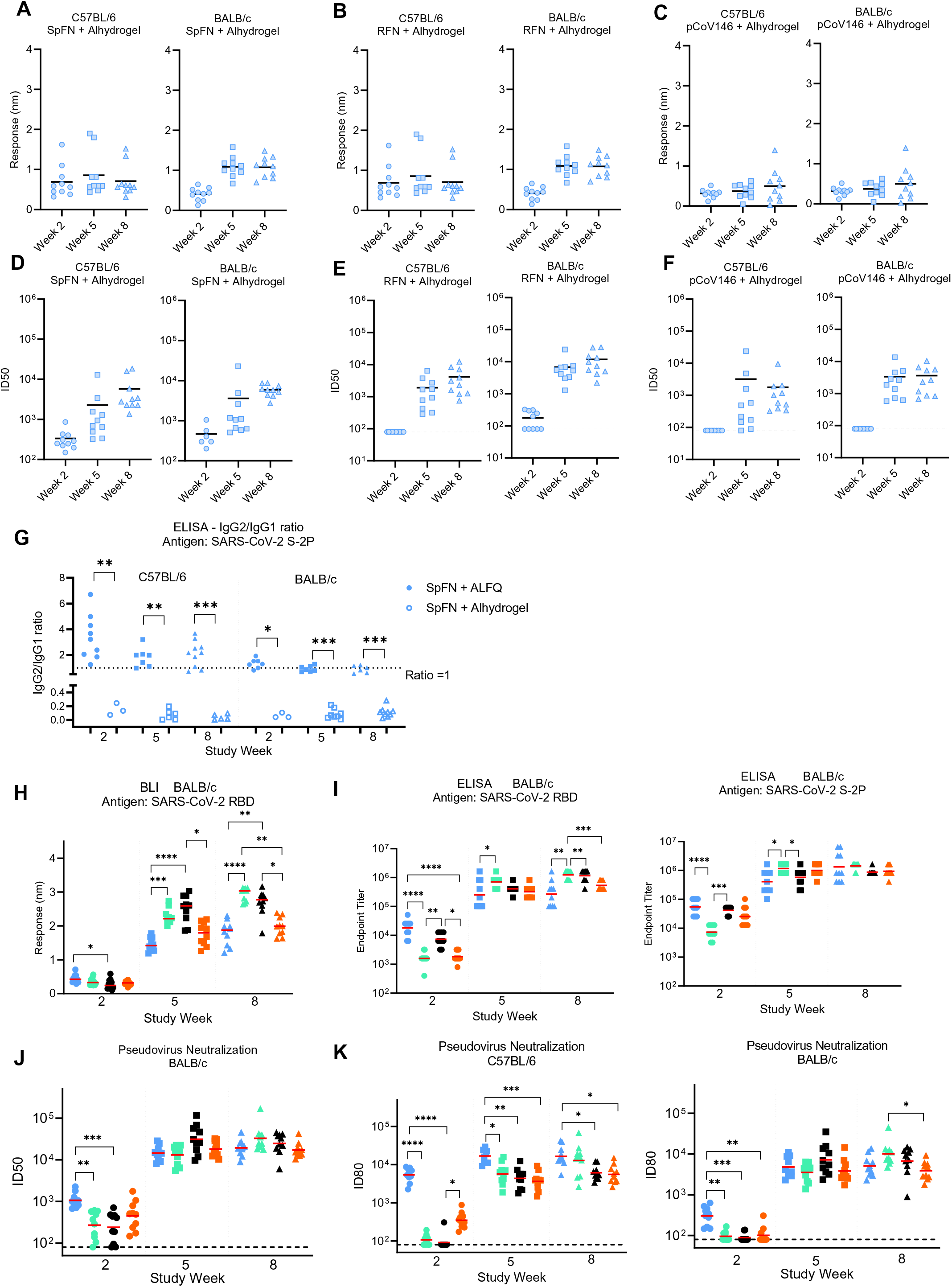
SARS-CoV-2 nanoparticle vaccine candidates elicit robust binding and pseudovirus neutralizing antibody responses in mice. Related to Figure 5 and 7. (A) Biolayer Interferometry binding analysis of C57BL/6 and BALB/c sera from mice immunized with SpFN + Alhydrogel® (B) RFN + Alhydrogel® and (C) pCoV146 + Alhydrogel® to SARS-CoV-2 RBD. Mean values are indicated by a horizontal line, n=10. (D) Pseudovirus neutralization (ID50 values) of C57BL/6 and BALB/c sera from mice immunized with SpFN + Alhydrogel® (E) RFN + Alhydrogel® and (F) pCoV146 + Alhydrogel®. Geometric mean values are indicated by a horizontal line, n=10. (G) ELISA analysis of antibody isotype usage following immunization with SpFN + ALFQ (solid shapes), or SpFN + Alhydrogel® (open shapes). Sera collected study week 2, 5, and 8 from immunized mice were added in quadruplicate serial dilutions to ELISA plates coated with S-2P protein. Duplicated wells were probed with anti-mouse-IgG1-HRP. Additional duplicates were probed with either anti-mouse-IgG2c-HRP or anti-mouse IgG2a-HRP for C57BL/6 and BALB/c mice, respectively. Data was interpolated to obtain the dilution factor at OD450 of 1 and plotted as ratios of IgG2/IgG1. A horizontal dotted line denotes a balanced 1:1 IgG2/IgG1 ratio. Isotype ratio values were compared between the two adjuvant groups at each timepoint for each mouse type using a Mann-Whitney unpaired two-tailed non-parametric test. (H) Biolayer interferometry analysis of BALB/c mouse sera binding to SARS-CoV-2 RBD at study weeks 2, 5 and 8. Mice were immunized with the four lead candidate vaccines SpFN (blue), RFN (green), pCoV146 (black) and pCOV111 (orange). Binding mean values are indicated by a horizontal line, n=10, sera responses at a given study week were compared for statistical differences using a Kruskal-Wallis test followed by a Dunn’s post-test. (I) ELISA analysis of BALB/c mice immune responses as indicated in (H). Binding geometric mean values of the endpoint titers are indicated by a horizontal line, n=10, sera responses at a given study week were compared for statistical differences using a Kruskal-Wallis test followed by a Dunn’s post-test. (J) Pseudovirus neutralization ID50 titers of BALB/c mice immunized as indicated in (H). Geometric mean values are indicated by a horizontal line, n=10, sera neutralization titers at a given study week for the four immunogens were compared for statistical differences using a Kruskal-Wallis test followed by a Dunn’s post-test. (K) Pseudovirus neutralization ID80 titers of C57BL/6 (left) and BALB/c mice (right) immunized as indicated in (H). Geometric mean values are indicated by a horizontal line, n=10, sera neutralization titers at a given study week for the four immunogens were compared for statistical differences using a Kruskal-Wallis test followed by a Dunn’s post-test. P values <0.0001 (****), <0.001 (***), <0.01 (**) or <0.05 (*).

**Figure S5.**
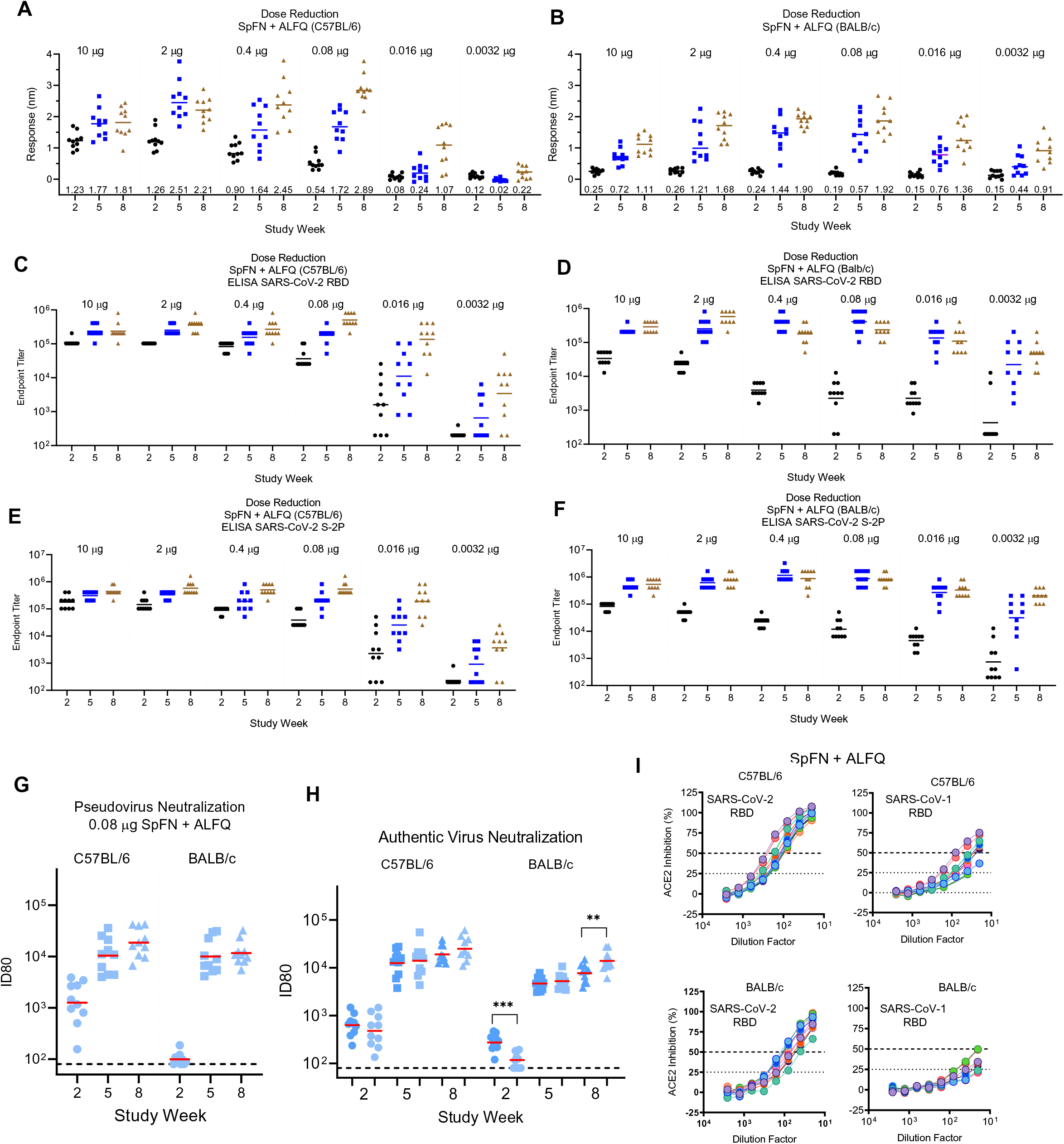
SARS-CoV-2 SpFN vaccine candidate elicits robust binding and neutralizing antibody responses at reduced doses in mice. Related to Figure 5 and 7. A) Biolayer interferometry analysis of C57BL/6 and (B) BALB/c mouse sera binding response to SARS-CoV-2 RBD following immunization with reducing doses of SpFN. (C, E) ELISA analysis of C57BL/6 and (D, F) BALB/c mouse sera binding response to SARS-CoV-2 RBD or S-2P following immunization with reducing doses of SpFN. (G) SARS-CoV-2 pseudovirus ID80 neutralization titers of mice immunized with 0.08 μg SpFN + ALFQ. (H) Authentic SARS-CoV-2 virus ID80 neutralization titers of mice immunized with 10 μg (blue) or 0.08 μg (light blue) SpFN + ALFQ. Geometric mean titers for each group and time point are indicated by a horizontal line, n =10. Neutralization titers for the two dose groups at each study time point were compared for statistically significant differences using a Mann-Whitney unpaired two-tailed non-parametric test. The two BALB/c time points that showed differences are indicated by bars. P values <0.001 (***), <0.01 (**). (I) Mouse sera from study week 10 was analyzed for hACE2 blocking capacity to SARS-CoV-2 RBD (left) or SARS-CoV-1 RBD using a biolayer interferometry assay format.

**Figure S6.**
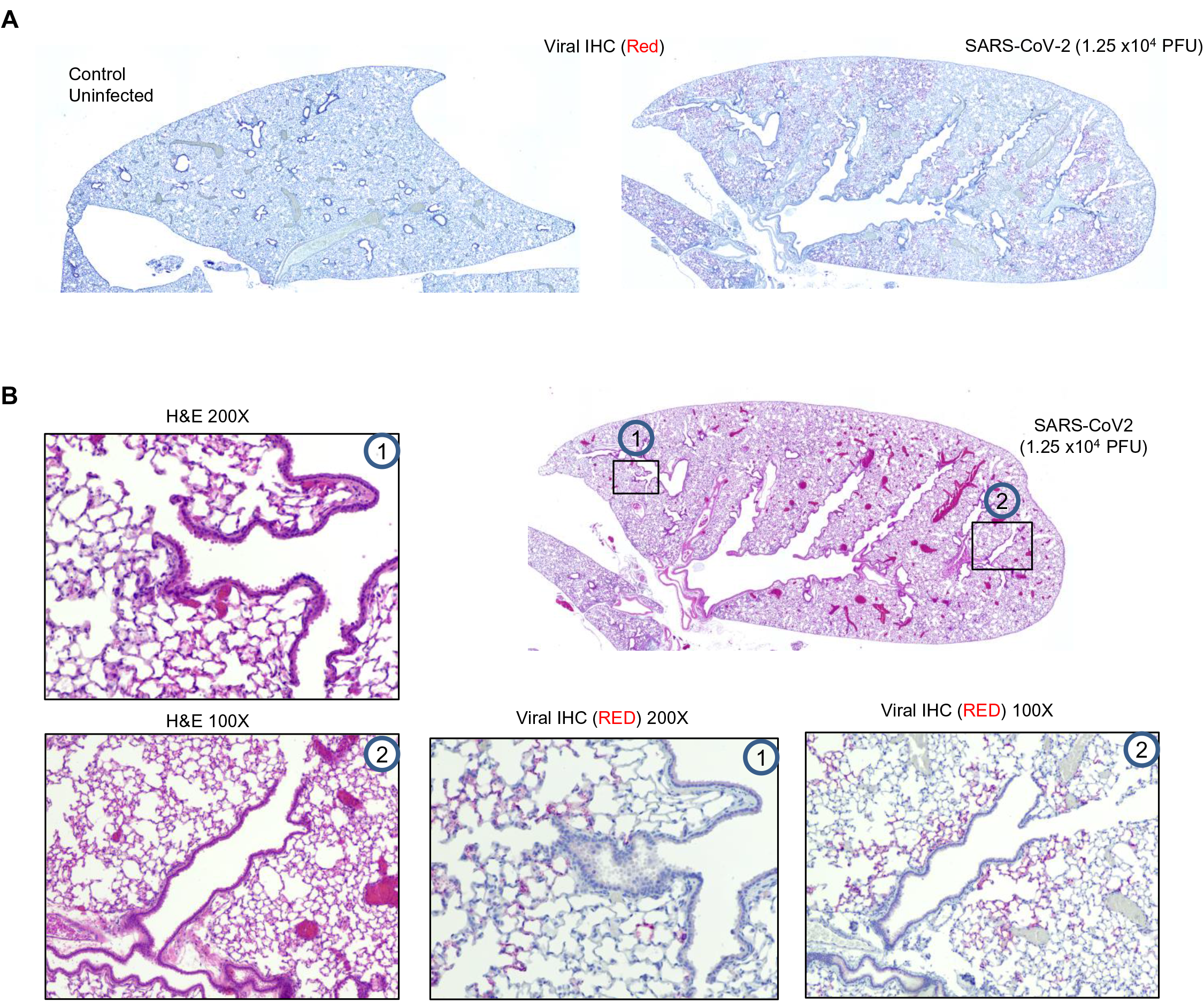
Histopathological analysis of SARS-CoV-2 infection in K18-ACE2 mice. Related to Figure 7. (A, B) Hematoxylin and eosin staining of lung sections from K18-hACE2 mice following intranasal infection with 1.25 × 10^4^ PFU SARS-CoV-2. Images show two magnifications. Images are representative of n = 10 per group.

**Table S1.** Spike-domain ferritin immunogens.

| Spike-Ferritin (all based on S-2P variant with Δfurin and PP) | | | |
| --- | --- | --- | --- |
| Construct ID | Description |  |  |
| pCoV1B-01 | S2P(1-1137)-del-4-Ferritin | Shortened ectodomain - no coiled coil (closest to flu HA pass off) | NL |
| pCoV1B-02 | S2P(1-1137)-del-6-Ferritin | Shortened ectodomain - no coiled coil (closest to flu HA pass off) | NL |
| pCoV1B-03 | S2P(1-1208)-del-Ferritin | Full ectodomain | NL |
| pCoV1B-04 | S2P(1-1208)-GCN4-Ferritin | Full ectodomain with GCN4 | NL |
| pCoV1B-05 | S2P(1-1154)-del-Ferritin | Shortened ectodomain with ending with a couple turns of coiled coil | NL |
| pCoV1B-06 | S2P(1-1158)op1-del-Ferritin | Optimized HR ending (end on glycan N1158) | NL |
| pCoV1B-07 | S2P(1-1158)op2-del-Ferritin | Optimized HR ending (Ile) (end on glycan N1158) | NL |
| pCoV1B-08 | S2P(1-1158)op1x2-del-Ferritin | Optimized HR ending (N1158 glycan removed, but exists on the repeated HR) | NL |
| pCoV1B-09 | S2P(1-1158)op2x2-del-Ferritin | Optimized HR ending (Ile) (N1158 glycan removed, but exists on the repeated HR) | NL |
| pCoV1B-10 | S2P(1-1158)op1-fGCN4-del-Ferritin | Optimized HR ending with GCN4 fused in register (no glycan N1158) | NL |
| pCoV1B-01-PL | PL-S2P(12-1137)-del-4-Ferritin | Shortened ectodomain - no coiled coil (closest to flu HA pass off) | PL |
| pCoV1B-02-PL | PL-S2P(12-1137)-del-6-Ferritin | Shortened ectodomain - no coiled coil (closest to flu HA pass off) | PL |
| pCoV1B-03-PL | PL-S2P(12-1208)-del-Ferritin | Full ectodomain | PL |
| pCoV1B-04-PL | PL-S2P(12-1208)-GCN4-Ferritin | Full ectodomain with GCN4 | PL |
| pCoV1B-05-PL | PL-S2P(12-1154)-del-Ferritin | Shortened ectodomain with ending with a couple turns of coiled coil | PL |
| pCoV-1B-06-PL (aka SpFN) | PL-S2P(12-1158)op1-del-Ferritin | Optimized HR ending (end on glycan N1158) | PL |
| pCoV1B-07-PL | PL-S2P(12-1158)op2-del-Ferritin | Optimized HR ending (Ile) (end on glycan N1158) | PL |
| pCoV1B-08-PL | PL-S2P(12-1158)op1x2-del-Ferritin | Optimized HR ending (N1158 glycan removed, but exists on the repeated HR) | PL |
| pCoV1B-09-PL | PL-S2P(12-1158)op2x2-del-Ferritin | Optimized HR ending (Ile) (N1158 glycan removed, but exists on the repeated HR) | PL |
| pCoV1B-10-PL | PL-S2P(12-1158)op1-fGCN4-del-Ferritin | Optimized HR ending with GCN4 fused in register (no glycan N1158) | PL |
| RBD-Ferritin |  |  |  |
| Construct ID | Description | Comment | Leader |
| pCoV03 | His8-3C-RBD(331-527)-Ferritin | N-terminal His8 with HRV-3C cleavage site, GSGGGG linker between RBD and Ferritin | PL |
| pCoV29 | His8-3C-RBD-3-Ferritin | SGG linker | PL |
| pCoV30 | His8-3C-RBD-3-del-Ferritin | SGG linker, $\Delta$ first 10 residues in ferritin, then DIEK changed to DIIK | PL |
| pCoV31 | His8-3C-RBD-6-del-Ferritin | P527G, $\Delta$ first 8 residues in ferritin, then SKDIEK changed to DIIK | PL |
| pCoV1A-01 | His8-3C-RBD-PPII-Ferritin | Extend distance between RBD and ferritin - using polyproline Helix | PL |
| pCoV1A-02 | His8-3C-RBD-alpha1-Ferritin | Extend distance between RBD and ferritin - using alpha Helix from bottom of S protein | PL |
| pCoV1A-03 | His8-3C-RBD-alpha2-Ferritin | Extend distance between RBD and ferritin- using alpha Helix from bottom of S protein | PL |
| pCoV1A-04 | His8-3C-RBD-GCN4-del-Ferritin | Extend distance between RBD and ferritin + stabilize ferritin - using GCN4 trimerization motif | PL |
| pCoV1A-05 | His8-3C-RBD-1141_1158op1-del-Ferritin | Extend distance between RBD and ferritin + stabilize ferritin - using semi-native trimerization motif | PL |
| pCoV1A-06 | His8-3C-RBD-1141_1158op1x2-del-Ferritin | Extend distance between RBD and ferritin + stabilize ferritin - using semi-native trimerization motif | PL |
| pCoV49 | His8-3C-RBD-F456N/K458T-Ferritin | RBD with indicated point mutations | PL |
| pCoV50 | His8-3C-RBD-L455R/Y449K/F490R-Ferritin | RBD with indicated point mutations | PL |
| pCoV51 | His8-3C-RBD-L455R-Ferritin | RBD with indicated point mutation | PL |
| pCoV52 | His8-3C-RBD-I468R-Ferritin | RBD with indicated point mutation | PL |
| pCoV53 | His8-3C-RBD-Y453R-Ferritin | RBD with indicated point mutation | PL |
| pCoV54 | His8-3C-RBD-L452R-Ferritin | RBD with indicated point mutation | PL |
| pCoV55 | His8-3C-RBD-L492R-Ferritin | RBD with indicated point mutation | PL |
| pCoV56 | His8-3C-RBD-F490R-Ferritin | RBD with indicated point mutation | PL |
| pCoV57 | His8-3C-RBD-F490A-Ferritin | RBD with indicated point mutation | PL |
| pCoV58 | His8-3C-RBD-L518N/L519K/H520S-Ferritin | RBD with indicated point mutations | PL |
| pCoV59 | His8-3C-RBD-L518R-Ferritin | RBD with indicated point mutation | PL |
| pCoV60 | His8-3C-RBD-V367T/L335N-Ferritin | RBD with indicated point mutations | PL |
| pCoV61 | His8-3C-RBD-T385N/L387T-Ferritin | RBD with indicated point mutations | PL |
| pCoV62 | His8-3C-RBD-V382R-Ferritin | RBD with indicated point mutation | PL |
| pCoV63 | His8-3C-RBD-F377R-Ferritin | RBD with indicated point mutation | PL |
| pCoV127 | His8-3C-RBD-F490A/L518N/L519K/H520S-Ferritin | RBD with indicated point mutations | PL |
| pCoV128 | His8-3C-RBD-F490A/L518R-Ferritin | RBD with indicated point mutations | PL |
| pCoV129 | His8-3C-RBD-L455R/Y449K/F490R/L518N/L519K/H520S-Ferritin | RBD with indicated point mutations | PL |
| pCoV130 | His8-3C-RBD-L455R/Y449K/F490R/L518R-Ferritin | RBD with indicated point mutations | PL |
| pCoV131 (aka RFN) | His8-3C-RBD-Y453R/L518N/L519K/H520S-Ferritin | RBD with indicated point mutations | PL |
| pCoV132 | His8-3C-RBD-Y453R/L518R-Ferritin | RBD with indicated point mutations | PL |

| RBD-NTD-Ferritin |  |  |  |
| --- | --- | --- | --- |
| Construct ID | Description |  |  |
| pCoV122 | His8-3C-RBD(331-527)-GSGGSG-NTD(12-303)-Ferritin | N-terminal His8 with HRV-3C cleavage site, GSGGSG linker between RBD and NTD, GSGGGG linker between NTD and Ferritin | PL |
| pCoV123 | His8-3C-RBD-F490R-NTD-Ferritin | RBD with indicated point mutation | PL |
| pCoV124 | His8-3C-RBD-F490A-NTD-Ferritin | RBD with indicated point mutation | PL |
| pCoV125 | His8-3C-RBD-L518N/L519K/H520S-NTD-Ferritin | RBD with indicated point mutations | PL |
| pCoV126 | His8-3C-RBD-L518R-NTD-Ferritin | RBD with indicated point mutation | PL |
| pCoV146 | His8-3C-RBD-Y453R-L518N/L519K/H520S-NTD-Ferritin | RBD with indicated point mutations | PL |
| pCoV147 | His8-3C-RBD-F490A-L518N/L519K/H520S-NTD-Ferritin | RBD with indicated point mutations | PL |

| S1-Ferritin |  |  |  |
| --- | --- | --- | --- |
| Construct ID | Description |  |  |
| pCoV68 | S1(12-678)-Ferritin | GSGGSG linker between S1 and Ferritin | PL |
| pCoV107 | S1(12-655)-Ferritin | 24 residues removed from the C-terminus | PL |
| pCoV108 | S1(12-655)-L611N/Q613T-Ferritin | 24 residues removed from the C-terminus, S1 with indicated point mutations | PL |
| pCoV109 | S1(12-696)-Ferritin | Extended the sequence to include a portion of S2 | PL |
| pCoV110 | S1(12-676)-G-S2(689-696)-Ferritin | Extended the sequence to include a portion of S2 with the indicated leader between the two regions | PL |
| pCoV111 | S1(12-676)-GG-S2(689-696)-Ferritin | Extended the sequence to include a portion of S2 with the indicated leader between the two regions | PL |
| pCoV112 | S1(12-676)-PG-S2(689-696)-Ferritin | Extended the sequence to include a portion of S2 with the indicated leader between the two regions | PL |
| pCoV113 | S1-Y312N/Q313Y/T314T-Ferritin | S1 with indicated point mutations | PL |
| pCoV114 | S1-I651N/A653S-Ferritin | S1 with indicated point mutations | PL |
| pCoV115 | S1-S316C/V595C-Ferritin | S1 with indicated point mutations | PL |
| pCoV116 | S1-V320C/S591C-Ferritin | S1 with indicated point mutations | PL |
| pCoV117 | S1-L560Q/F562H-Ferritin | S1 with indicated point mutations | PL |
| pCoV118 | S1-F562N/Q564T-Ferritin | S1 with indicated point mutations | PL |
| pCoV119 | S1-F490R-Ferritin | S1 with indicated point mutation | PL |
| pCoV120 | S1-F490A-Ferritin | S1 with indicated point mutation | PL |
| pCoV02 | S1(16-678)-Ferritin | 4 residues removed from N-terminus | PL |
| pCoV67 | His8-3C-S1-Ferritin | His8 and HRV-3C cleavage site added to N-terminus | PL |

**Table S2.** Negative-stain Electron Microscopy Data Collection and Refinement.

| Protein | SpFN_1B-06-PL | RFN_131 | pCoV146 | pCoV111 | pCoV1B-05 |
| --- | --- | --- | --- | --- | --- |
| Immunogen Fused | Spike (S2P) | RBD | RBD-NTD | S1 | Spike (S2P) |
| EMPIAR Code | XXXXXX | XXXXXX | XXXXXX | XXXXXX | XXXXXX |
| EMDB Code | XXXX | XXXX | XXXX | XXXX | XXXX |
| **Data Collection** |  |  |  |  |  |
| Microscope | Tecnai T20 | Tecnai T20 | Tecnai T20 | Tecnai T20 | Talos L120C |
| Voltage (kV) | 200 kV | 200 kV | 200 kV | 200 kV | 120 kV |
| Camera | Eagle 4K | Eagle 4K | Eagle 4K | Eagle 4K | Ceta |
| Software | SerialEM | SerialEM | SerialEM | SerialEM | EPU |
| Pixel Size (Å/pix) | 2.195 | 2.195 | 2.195 | 2.195 | 2.542 |
| Underfocus range | 0.7-1.3 | 0.8-1.3 | 0.6-1.5 | 0.8-1.6 | 0.5-0.9 |
| **Image Processing** |  |  |  |  |  |
| Software | RELION | RELION | RELION | RELION | RELION |
|  | 3.0.8 | 3.0.8 | 3.0.8 | 3.0.8 | 3.1.1 |
| # Particle Images | 11502 | 3383 | 832 | 2121 | 2143 |
| Pixel Size (Å/pixel) | 4.39 | 4.39 | 4.39 | 4.39 | 5.084 |
| Box Size (pixels) | 160 | 160 | 160 | 160 | 200 |
| Symmetry (3D) | O | O | O | O | -- |
| Initial Lowpass (Å) (RELION) | 100 | 80 | 100 | 100 | -- |
| High-res Limit (Å) (cisTEM) | -- | -- | -- | -- | -- |
| Resolution (Å) | 25 | 21 | 30 | 30 | -- |

**Table S3.** Mouse immunogenicity study immunogens, adjuvants, and mouse type.

| pCOV no. | Immunogen design category,<br>Study design | C57BL/6<br>ALFQ | Balb/c<br>ALFQ | C57BL/6<br>Alhydrogel | Balb/c<br>Alhydrogel |
| --- | --- | --- | --- | --- | --- |
| 1B-05 | S-Trimer-Ferritin | X | X |  |  |
| 1B-06-PL | S-Trimer-Ferritin | X | X | X | X |
| **RBD-Ferritin constructs** |  |  |  |  |  |
| pCOV no. | Immunogen design category,<br>Study design | C57BL/6<br>ALFQ | Balb/c<br>ALFQ | C57BL/6<br>Alhydrogel | Balb/c<br>Alhydrogel |
| 50 | RBD-Ferritin |  | X |  |  |
| 58 | RBD-Ferritin | X | X | X | X |
| 59 | RBD-Ferritin |  | X |  |  |
| 127 | RBD(57+58)-Ferritin | X | X | X | X |
| 129 | RBD(50+58)-Ferritin | X | X | X | X |
| 130 | RBD(50+59)-Ferritin |  | X |  |  |
| 131 | RBD(53+58)-Ferritin | X | X | X | X |
| **S1-Ferritin constructs** |  |  |  |  |  |
| pCOV no. |  | C57BL/6<br>ALFQ | Balb/c<br>ALFQ | C57BL/6<br>Alhydrogel | Balb/c<br>Alhydrogel |
| 111 | S1-Ferritin | X | X |  |  |
| **RBD-NTD-Ferritin constructs** |  |  |  |  |  |
| pCOV no. | Immunogen design category,<br>Study design | C57BL/6<br>ALFQ | Balb/c<br>ALFQ | C57BL/6<br>Alhydrogel | Balb/c<br>Alhydrogel |
| 122 | RBD-NTD-Ferritin | X | X |  |  |
| 125 | RBD(58)-NTD-Ferritin |  | X |  | X |
| 146 | RBD(53+58)-NTD-Ferritin | X | X | X | X |
| 147 | RBD(57+58)-NTD-Ferritin | X |  |  |  |

**Table S4.** Animal immunogenicity SARS-CoV-2 pseudovirus neutralization ID50 and ID80. Numbers shown are the ID50/ID80 geometric mean titers for a group, with study week 2, 5, and 8 shown in vertical order.

| pCOV no. | Immunogen design category,<br>Study design | C57BL/6<br>ALFQ | Balb/c<br>ALFQ | C57BL/6<br>Alhydrogel | Balb/c<br>Alhydrogel |
| --- | --- | --- | --- | --- | --- |
| 1B-05 | S-Trimer-Ferritin (x 2<br>groups) | 702/189<br>8,709/2,346<br>13,076/5,647 | 115/<80<br>3,934/716<br>5,546/1,447 |  |  |
| 1B-06-PL | S-Trimer-Ferritin | 14,976/5,396<br>41,237/16,8184<br>7,323/16,52 | 1,152/355<br>16,816/6,662<br>25,062/6,540 |  |  |
| **RBD-Ferritin constructs** |  |  |  |  |  |
| pCOV no. | Immunogen design category,<br>Study design | C57BL/6<br>ALFQ | Balb/c<br>ALFQ | C57BL/6<br>Alhydrogel | Balb/c<br>Alhydrogel |
| 50 | RBD-Ferritin |  | X |  |  |
| 58 | RBD-Ferritin | 577/238<br>11,224/2,793<br>31,562/10,09 | 353/123<br>13,466/3,802<br>25,340/7,692 | 293/211<br>1,734/688<br>5,097/1261 | 232/<80<br>4,836/1,086<br>9,439/2,569 |
| 59 | RBD-Ferritin |  | X |  |  |
| 127 | RBD(57+58)-Ferritin | X | X | X | X |
| 129 | RBD(50+58)-Ferritin | X | X | X | X |
| 130 | RBD(50+59)-Ferritin |  | X |  |  |
| 131 | RBD(53+58)-Ferritin | 358/107<br>15,950/5,667<br>38,110/12,824 | 270/95<br>13,090/3,539<br>32,969/10,079 | 682/163<br>1,181/403<br>2,845/529 | 119/<40<br>182/103<br>240/99 |
| **S1-Ferritin constructs** |  |  |  |  |  |
| pCOV no. |  | C57BL/6<br>ALFQ | Balb/c<br>ALFQ | C57BL/6<br>Alhydrogel | Balb/c<br>Alhydrogel |
| 111 | S1-Ferritin | 1,770/350<br>14,893/3,636<br>19,157/5,564 | 450/172<br>18,112/3,846<br>17,108/3,886 |  |  |
| **RBD-NTD-Ferritin constructs** |  |  |  |  |  |
| pCOV no. | Immunogen design category,<br>Study design | C57BL/6<br>ALFQ | Balb/c<br>ALFQ | C57BL/6<br>Alhydrogel | Balb/c<br>Alhydrogel |
| 122 | RBD-NTD-Ferritin | X | X |  |  |
| 125 | RBD(58)-NTD-Ferritin |  | X |  | X |
| 146 | RBD(53+58)-NTD-Ferritin | 230/91<br>16,678/4,356<br>20,107/6,126 | 240/89<br>31,252/7,190<br>24,854/6,744 | <80/<80<br>667/460<br>940/289 | 662/<80<br>2,087/537<br>2,417/701 |
| 147 | RBD(57+58)-NTD-Ferritin | X |  |  |  |

## Materials and Methods

### Immunogen Modeling and Design

Following release of the SARS-CoV-2 sequence on Jan 10^th^ 2020, initial RBD-Ferritin and S1-Ferritin immunogens were designed (Table S1). Subsequent iterative immunogen design and optimization utilized atomic models of the SARS-CoV-2 RBD molecule (Joyce et al., 2020), or the SARS-2 S trimer structure PDB ID: 6VXX, and PDB ID: 3BVE for the *Helicobacter pylori* Ferritin, and PDB ID: 4LQH for the bullfrog linker sequence. Pymol (Schrödinger) was used to generate the ferritin 24-subunit particle, and a map created in UCSF Chimera (Pettersen et al., 2004) was supplied to cisTEM (Grant et al., 2018) “align_symmetry” to align the ferritin particle to an octahedral symmetry convention. This was supplied to “phenix.map_symmetry” to generate a symmetry file and PDB file, for octahedral (for monomer-fusions) and D4 (for trimer-fusions) symmetry. S-domain ferritin nanoparticle fusions were modelled using Pymol and Coot (Emsley et al., 2010) and expanded using “phenix.apply_ncs” (Liebschner et al., 2019). Visual analysis and figure generation was conducted using ChimeraX and PyMOL.

RBD-Ferritin designs were generated by assessment of the hydrophobic surface of the SARS-CoV-2 RBD surface and determining surface accessible mutations that reduced the hydrophobic surface. S1-Ferritin designs were creating using the PDB ID: 6VXX and including a short region of the S2 domain, which interacts with S1. Spike-Ferritin designs were created by modeling the coiled-coil region between S residues 1140 and 1158 and increasing the coil-coil interaction either by mutagenesis, or by increasing the length of the interaction region. RBD-NTD-Ferritin designs utilized RBD constructs with improved properties in the context of RBD-Ferritin, which were fused to the N-terminus of NTD (12 - 303)-Ferritin by a short 6 amino-acid linker.

### DNA plasmid construction and preparation

SARS-CoV-2 S-domain ferritin constructs were derived from the Wuhan-Hu-1 strain genome sequence (GenBank MN9089473), to include the following domains: RBD subunit (residues 331 - 527), NTD subunit (residues 12 - 303), RBD subunit linked to NTD subunit (residues 331 - 527 linked to residues 12 - 303 with a short GSG linker), S1 domain (residues 12 - 696) and S ectodomain (residues 12 – 1158). Constructs were modified to incorporate a N-terminal hexa-histadine tag (his) for purification of the RBD-Ferritin and RBD-NTD-Ferritin constructs.

An S-2P construct was used as a template to generate the set of Spike ferritin nanoparticles. Subsequent designs involving small deletions, additions and point mutations were generated using a modified QuikChange site-directed mutagenesis protocol (Agilent). RBD-ferritin and S1-ferritin constructs were synthesized by GenScript. For some of the RBD-ferritin constructs, gene segments (gBlocks) were synthesized by Integrated DNA Technologies to encode various linker regions between RBD and ferritin. Gene segments were stitched together with RBD- and ferritin-encoding PCR products using overlap extension PCR and were re-subcloned into the CMVR vector. The His-tagged SARS-CoV-2 RBD molecule was generated by amplifying the RBD domain from the RBD-Ferritin plasmid while encoding the 3’ purification tag and subcloned into the CMVR vector. The NTD protein subunit was generated in a similar manner, by amplifying the NTD domain from the S1-Ferritin construct. For expression of S, RBD, and NTD proteins, the S protein domains were cloned into the CMVR expression plasmid using the NotI/BamHI restriction sites. Constructs including the N-terminal region of the S protein included the native leader sequence; for constructs not including this region we utilized a prolactin leader (PL) sequence (Boyington et al., 2016).

Plasmid DNA generated by subcloning (restriction digest and ligation) was amplified in and isolated from E. coli Top10 cells. The constructs resulting from site-directed mutagenesis were either amplified in and isolated from E. coli Stbl3 or Top10 cells. Large-scale DNA isolation was performed using either endo free Maxiprep, Megaprep or Gigaprep kits (Qiagen).

### Immunogen expression and purification

All expression vectors were transiently transfected into Expi293F cells (Thermo Fisher Scientific) using ExpiFectamine 293 transfection reagent (Thermo Fisher Scientific). Cells were grown in polycarbonate baffled shaker flasks at 34°C or 37°C and 8% CO_2_ at 120 rpm. Cells were harvested 5-6 days post-transfection via centrifugation at 3,500 × g for 30 minutes. Culture supernatants were filtered with a 0.22-µm filter and stored at 4 °C prior to purification.

His-tagged proteins were purified using Ni-NTA affinity chromatography, while untagged proteins were purified with GNA lectin affinity chromatography. Briefly, 25 mL GNA-lectin resin (VectorLabs) was used to purify untagged protein from 1L of expression supernatant. GNA resin was equilibrated with 10 column volumes (CV) of phosphate buffered saline (PBS) (pH 7.4) followed by supernatant loading twice at 4 °C. Unbound protein was removed by washing with 20 CV of PBS buffer. Bound protein was eluted with 250mM methyl-α -D mannopyranoside in PBS buffer (pH 7.4). His-tagged proteins were purified using Ni-NTA affinity chromatography. 1 mL Ni-NTA resin (Thermo Scientific) was used to purify protein from 1L of expression supernatant. Ni-NTA resin was equilibrated with 5 CV of phosphate buffered saline (PBS) (pH 7.4) followed by supernatant loading 2x at 4 °C. Unbound protein was removed by washing with 200 CV of PBS, followed by 50 CV 10mM imidazole in PBS. Bound protein was eluted with 220mM imidazole in PBS. For all proteins, purification purity was assessed by SDS-PAGE. RBD-Ferritin nanoparticle constructs had a propensity to form soluble or insoluble aggregates which affected the ability to concentrate the samples. Addition of 1mM EDTA and 5% glycerol to the NiNTA purified material, prior to SEC or other concentration steps, mitigated the aggregation issue, and increased the nanoparticle formation as judged by SEC, and confirmed by neg-EM. All proteins were further purified by size-exclusion chromatography using a 16/60 Superdex-200 purification column. Purification purity for all the proteins was assessed by SDS-PAGE. Removal of the His-tags for SARS2-CoV-2 S-2P and RBD for use in ELISA were carried out using HRV-3C protease. Endotoxin levels for ferritin nanoparticle immunogens were assessed (Endosafe® nexgen-PTS, Charles River Laboratories) and 5 % v/v glycerol was added prior to filter-sterilization with a 0.22-µm filter, flash-freezing in liquid nitrogen, and storage at −80 °C. Ferritin nanoparticle formation was assessed by dynamic light scattering (DLS) by determining the hydrodynamic diameter at 25 °C using a Malvern Zetasizer Nano S (Malvern, Worcestershire, UK) equipped with a 633-nm laser.

For the antibodies, plasmids encoding heavy and light chains of antibodies (CR3022, and SR1-SR5) were co-transfected into Expi293F cells (ThermoFisher) according to the manufacturer’s instructions for expression of antibodies. After 5 days, antibodies were purified from cleared culture supernatants with Protein A agarose (ThermoFisher) using standard procedures, buffer was exchanged to PBS by dialysis, and antibody concentration was quantified using calculated extinction coefficient and A280 measurements.

### Negative-stain Electron Microscopy

Purified proteins were deposited at 0.02-0.08 mg/ml on carbon-coated copper grids and stained with 0.75% uranyl formate. Grids were imaged using a FEI T20 operating at 200 kV with an Eagle 4K CCD using SerialEM or using a Thermo Scientific Talos L120C operating at 120 kV with Thermo Scientific Ceta using EPU. All image processing steps were done using RELION 3.0.8, RELION 3.1.1, and/or cisTEM-1.0.0-beta. Particles were picked either manually or using templates generated from manually picked 2D class averages. CTF estimation was done with CTFFIND 4.1.13 and used for 2D classification. 3D reconstructions were generated using an initial reference generated from a corresponding synthetic atomic model with a low pass filter of 80-100 angstroms to remove distinguishable features or from a similar construct also low pass filtered to 80-100 angstroms. For all 3D reconstructions, O symmetry was enforced, and no explicit mask was used. Visual analysis and figure generation was conducted using Chimera and ChimeraX.

### Dynamic Light Scattering

Spike-domain ferritin nanoparticle hydrodynamic diameter was determined at 25°C using a Malvern Zetasizer Nano S (Malvern, Worcestershire, UK) equipped with a 633-nm laser. Samples were assessed accounting for the viscosity of their respective buffers.

### Octet Biolayer Interferometry binding and ACE2 inhibition assays

All biosensors were hydrated in PBS prior to use. All assay steps were performed at 30°C with agitation set at 1,000 rpm in the Octet RED96 instrument (FortéBio). Biosensors were equilibrated in assay buffer (PBS) for 15 seconds before loading of IgG antibodies (30 µg/ml diluted in PBS). SARS-COV-2 antibodies were immobilized onto AHC biosensors (FortéBio) for 100 seconds, followed by a brief baseline in assay buffer for 15 s. Immobilized antibodies were then dipped in various antigens for 100-200 s followed by dissociation for 20-100 s.

Mouse sera binding to the SARS-CoV-2 RBD, VoC RBDs, or SARS-CoV-1 RBD were carried out as follows. HIS1K biosensors(FortéBio) were equilibrated in assay buffer for 15 s before loading of His-tagged RBD (30 μg/ml diluted in PBS) for 120 seconds. After briefly dipping in assay buffer (15 seconds in PBS), the biosensors were dipped in mouse sera (100-fold dilution) for 180 seconds followed by dissociation for 60 seconds.

SARS-CoV-2 and SARS-CoV-1 RBD hACE2 inhibition assays were carried out as follows. SARS-CoV-2 or SARS-CoV-1 RBD (30 μg/ml diluted in PBS) was immobilized on HIS1K biosensors (FortéBio) for 180 seconds followed by baseline equilibration for 30 s. Serum was allowed to occur for 180 s followed by baseline equilibration (30 s). ACE2 protein (30 μg/ml) was the allowed to bind for 120 s. Percent inhibition (PI) of RBD binding to ACE2 by serum was determined using the equation: PI = 100 − [(ACE2 binding in the presence of mouse serum)) ⁄ (mouse serum binding in the absence of competitor mAb)] × 100.

### Mouse immunization

All research in this study involving animals was conducted in compliance with the Animal Welfare Act, and other federal statutes and regulations relating to animals and experiments involving animals and adhered to the principles stated in the Guide for the Care and Use of Laboratory Animals, NRC Publication, 1996 edition. The research protocol was approved by the Institutional Animal Care and Use Committee of WRAIR. BALB/c and C57BL/6 mice were obtained from Jackson Laboratories (Bar Harbor, ME). Mice were housed in the animal facility of WRAIR and cared for in accordance with local, state, federal, and institutional policies in a National Institutes of Health American Association for Accreditation of Laboratory Animal Care-accredited facility.

C57BL/6 or BALB/c mice (n=10/group) were immunized intramuscularly with 10 µg of immunogen (unless stated) adjuvanted with either ALFQ or Alhydrogel® in alternating caudal thigh muscles three times, at 3-week intervals; blood was collected 2 weeks before the first immunization, the day of the first immunization, and 2 weeks following each immunization, and at week 10 (Table S1). For immunogen SpFN_1B-06-PL, mice were immunized with reduced doses of protein adjuvanted with ALFQ with immunization schedule, site of injections, and timing of bleeds as described. Mice were randomly assigned to experimental groups and were not pre-screened or selected based on size or other gross physical characteristics. Serum was stored at 4°C or −80°C until analysis. Antibody responses were analyzed by Octet Biolayer Interferometry, enzyme-linked immunosorbent assay (ELISA), pseudovirus neutralization assay, and live-virus neutralization assay.

### Immunogen-Adjuvant preparation

Purified research grade nanoparticle immunogens were formulated in PBS with 5% glycerol at 1 mg/ml and subsequently diluted with dPBS (Quality Biological) to provide 10 μg or lower amount per 50 μl dose upon mixing with adjuvant. ALFQ (1.5X) liposomes, containing 600 µg/mL 3D-PHAD and 300 μg ug/mL QS-21, were gently mixed by slow speed vortex prior to use. Antigen was added to the ALFQ, vortexed at a slow speed for 1 minute, mixed on a roller for 15 minutes, and stored at 4°C for 1 h prior to immunization. Spike-Ferritin nanoparticle immunogens were formulated with ALFQ to contain 20 μg 3D-PHAD and 10 μg QS21 per 50 μl dose. Alhydrogel® stock (10 mg/ml aluminum (GMP grade; Brenntag)) was diluted to 900 µg/mL (1.5X) with DPBS and gently mixed. Appropriate volume and concentration of antigen was added to the diluted Alhydrogel® before being vortexed at low speed for 1 min, mixed on a roller for 15 minutes, and stored at 4°C for at least 1 h prior to immunization. Spike-Ferritin nanoparticle immunogens were adsorbed to Alhydrogel® aluminum hydroxide at 30 μg aluminum per 50 μl dose.

### Enzyme Linked Immunosorbent Assay (ELISA)

96-well Immulon “U” Bottom plates were coated with 1 μg/mL of RBD or S protein (S-2P) antigen in PBS, pH 7.4. Plates were incubated at 4°C overnight and blocked with blocking buffer (Dulbecco’s PBS containing 0.5% milk and 0.1% Tween 20, pH 7.4, at room temperature (RT) for 2 h. Individual serum samples were serially diluted 2-fold in blocking buffer and added to triplicate wells and the plates were incubated for 1 hour (h) at RT. Horseradish peroxidase (HRP)-conjugated sheep anti-mouse IgG, gamma chain specific (The Binding Site) was added and incubated at RT for an hour, followed by the addition of 2,2’-Azinobis [3-ethylbenzothiazoline-6-sulfonic acid]-diammonium salt (ABTS) HRP substrate (KPL) for 1 h at RT. The reaction was stopped by the addition of 1% SDS per well and the absorbance (A) was measured at 405 nm (A405) using an ELISA reader Spectramax (Molecular Devices, San Jose, CA)) within 30 min of stopping the reaction.. Antibody positive (anti-RBD mouse mAb; BEI resources) and negative controls were included on each plate. The results are expressed as end point titers, defined as the reciprocal dilution that gives an absorbance value that equals twice the background value (antigen-coated wells that did not contain the test sera, but had all other components added).

Mouse isotype ELISA were performed using a similar approach as above, but with the following differences. Only Spike protein (S-2P) was used to coat the wells. The plates were blocked with PBS containing 0.2% bovine serum albumin (BSA), pH 7.4 for 30 minutes. The mouse serum samples were serially diluted in duplicates either 3- or 4-fold in PBS containing 0.2% BSA and 0.05% Tween 20, pH7.4. The secondary antibodies were HRP-conjugated AffiniPure Goat Anti-Mouse antibodies from Jackson ImmunoResearch specific for either Fcγ subclass 1, Fcγ subclass 2a, or Fcγ subclass 2c. The secondary antibodies were incubated for 30 minutes. TMB (3,3’,5,5’-Tetramethylbenzidine) substrate (Thermo) was added and the plates were incubated at RT for 5-10 minutes to allow color development. Stop solution (Thermo) was added and the A405 was measured using a VersaMax microplate reader (Molecular Devices). A titration curve of serum concentration versus A450 was created. The titration curves were interpolated to determine the dilution factor where A450=1.0 for each mouse sera sample and IgG subclass, and the resulting values were used to calculate the IgG1/IgG2a ratio (for BALB/c mice) or IgG1/IgG2c ratio (for C57BL/6 mice). In the animal groups immunized with SpFN_1B-06-PL with the adjuvant Alhydrogel®, many of the mouse IgG usage ratio could not be calculated due to insufficient signal for either IgG2a or IgG2c in the mouse sera.

### SARS-CoV-2 and SARS-CoV-1 pseudovirus neutralization assay

The S expression plasmid sequences for SARS-CoV-2 (Wuhan1, B.1.1.7, and B.1.351) and SARS-CoV were codon optimized and modified to remove an 18 amino acid endoplasmic reticulum retention signal in the cytoplasmic tail in the case of SARS-CoV-2, and a 28 amino acid deletion in the cytoplasmic tail in the case of SARS-CoV. This allowed increased S incorporation into pseudovirions (PSV) and thereby enhance infectivity. Virions pseudotyped with the vesicular stomatitis virus (VSV) G protein were used as a non-specific control. SARS-CoV-2 pseudovirions (PSV) were produced by co-transfection of HEK293T/17 cells with a SARS-CoV-2 S plasmid (pcDNA3.4) and an HIV-1 NL4-3 luciferase reporter plasmid (The reagent was obtained through the NIH HIV Reagent Program, Division of AIDS, NIAID, NIH: Human Immunodeficiency Virus 1 (HIV-1) NL4-3 ΔEnv Vpr Luciferase Reporter Vector (pNL4-3.Luc.R-E-), ARP-3418, contributed by Dr. Nathaniel Landau and Aaron Diamond). The SARS-CoV-2 S expression plasmid sequence was derived from the Wuhan seafood market pneumonia virus isolate Wuhan-Hu-1, complete genome (GenBank accession MN908947), and the SARS-CoV-1 expression plasmid was derived from the Urbani S sequence.

Infectivity and neutralization titers were determined using ACE2-expressing HEK293 target cells (Integral Molecular) in a semi-automated assay format using robotic liquid handling (Biomek NXp Beckman Coulter). Test sera were diluted 1:40 in growth medium and serially diluted, then 25 µL/well was added to a white 96-well plate. An equal volume of diluted SARS-CoV-2 PSV was added to each well and plates were incubated for 1 hour at 37°C. Target cells were added to each well (40,000 cells/ well) and plates were incubated for an additional 48 hours. RLUs were measured with the EnVision Multimode Plate Reader (Perkin Elmer, Waltham, MA) using the Bright-Glo Luciferase Assay System (Promega Corporation, Madison, WI). Neutralization dose–response curves were fitted by nonlinear regression with a five-parameter curve fit using the LabKey Server® (Piehler et al., 2011), and the final titers are reported as the reciprocal of the dilution of serum necessary to achieve 50% neutralization (ID50, 50% inhibitory dilution) and 80% neutralization (ID80, 80% inhibitory dilution). Assay equivalency for SARS-CoV-2 was established by participation in the SARS-CoV-2 Neutralizing Assay Concordance Survey (SNACS) run by the Virology Quality Assurance Program and External Quality Assurance Program Oversite Laboratory (EQAPOL) at the Duke Human Vaccine Institute, sponsored through programs supported by the National Institute of Allergy and Infectious Diseases, Division of AIDS.

### SARS-CoV-2 authentic virus neutralization assay

The neutralization assay has been described in detail previously (Case et al., 2020). Briefly, SARS-CoV-2 strain 2019-nCoV/USA_WA1/2020 was obtained from the Centers for Disease Control and Prevention. Virus was passaged once in Vero CCL81 cells (ATCC) and titrated by focus-forming assay on Vero E6 cells. Mouse sera were serially diluted and incubated with 100 focus-forming units of SARS-CoV-2 for 1 h at 37°C. Serum-virus mixtures were then added to Vero E6 cells in 96-well plates and incubated for 1 h at 37°C. Cells were overlayed with 1% (w/v) methylcellulose in MEM. After 30 h, cells were fixed with 4% PFA in PBS for 20 minutes at room temperature then washed and stained overnight at 4°C with 1 µg/ml of CR3022 (ter Meulen et al., 2006; Tian et al., 2020) in PBS supplemented with 0.1% saponin and 0.1% bovine serum albumin. Cells were subsequently stained with HRP-conjugated goat anti-human IgG for 2 h at room temperature. SARS-CoV-2-infected cell foci were visualized with TrueBlue peroxidase substrate (KPL) and quantified using ImmunoSpot® microanalyzer (Cellular Technologies, Shaker Heights, OH). Neutralization curves were generated using Prism software (GraphPad Prism 8.0).

### Protection experiments in K18-hACE2 transgenic mice

All research in this study involving animals was conducted in compliance with the Animal Welfare Act, and other federal statutes and regulations relating to animals and experiments involving animals and adhered to the principles stated in the Guide for the Care and Use of Laboratory Animals, NRC Publication, 1996 edition. The research protocol was approved by the Institutional Animal Care and Use Committee of the Trudeau Institure. K18-hACE2 transgenic mice were obtained from Jackson Laboratories (Bar Harbor, ME). Mice were housed in the animal facility of the Trudeau Institute and cared for in accordance with local, state, federal, and institutional policies in a National Institutes of Health American Association for Accreditation of Laboratory Animal Care-accredited facility.

To determine an appropriate challenge viral stock amount and establish the K18-hACE2 SARS-CoV-2 mouse challenge model, five viral doses were used to inoculate the K18-hACE2 mice. Each study group was composed of 10 hACE2 K18 Tg mice (5 males and 5 females). Mice were infected on study day 0 doses ranging from 5 × 10^2^ to 1 × 10^5^ PFU of SARS-CoV-2 USA-WA1/2020 administered via intranasal instillation. All mice were monitored for clinical symptoms and body weight twice daily, every 12 hours, from study day 0 to study day 14. Mice were euthanized if they displayed any signs of pain or distress as indicated by the failure to move after stimulated or inappetence, or if mice have greater than 25% weight loss compared to their study day 0 body weight. Hematoxylin and eosin staining of lung sections following infection with 1.25 × 10^4^ PFU compared to control uninfected mouse lung sections are shown in Figure S6.

For the passive immunization study, on day −1, K18-hACE2 mice were injected intravenously with purified IgG from C57BL/6 vaccinated mice. On study day 0, all mice were inoculated with 1.25×10^4^ PFU of SARS-CoV-2 USA-WA1/2020 via intranasal instillation. All mice were monitored for clinical symptoms and body weight twice daily, every 12 hours, from study day 0 to study day 14. Mice were euthanized if they displayed any signs of pain or distress as indicated by the failure to move after stimulated or inappetence, or if mice have greater than 25% weight loss compared to their study day 0 body weight.

### Data Analysis

Data analyses used GraphPad (San Diego, CA) Prism software and statistical tests as described for individual experiments.

